# Melanopsin activates divergent phototransduction pathways in ipRGC subtypes

**DOI:** 10.1101/2022.06.12.495838

**Authors:** Ely Contreras, Takuma Sonoda, Lutz Birnbaumer, Tiffany M. Schmidt

**Affiliations:** Department of Neurobiology, Northwestern University, Evanston, IL, USA; Northwestern University Interdisciplinary Biological Sciences Program, Northwestern University, Evanston, IL; Northwestern University Interdepartmental Neuroscience Program, Northwestern University, Chicago, IL; Laboratory of Signal Transduction, National Institute of Environmental Health Sciences, Research Triangle Park, North Carolina 27709 and Institute of Biomedical Research (BIOMED), Catholic University of Argentina, C1107AZZ Buenos Aires, Argentina; Department of Ophthalmology, Feinberg School of Medicine, Chicago, IL, USA

## Abstract

Melanopsin signaling within ipRGC subtypes impacts a broad range of behaviors from circadian photoentrainment to conscious visual perception. Yet, how melanopsin phototransduction within M1-M6 ipRGC subtypes impacts cellular signaling to drive diverse behaviors is still largely unresolved. The identity of the phototransduction channels in each subtype is key to understanding this central question but has remained controversial. In this study, we resolve two opposing models of M4 phototransduction, demonstrating that hyperpolarization-activated cyclic nucleotide-gated (HCN) channels are dispensable for this process and providing support for a model involving melanopsin-dependent potassium channel closure and canonical transient receptor potential (TRPC) channel opening. Surprisingly, we find that HCN channels are likewise dispensable for M2 phototransduction, contradicting the current model. We instead show that in M2 phototransduction, TRPC channels act in conjunction with T-type voltage-gated calcium channels, a novel melanopsin phototransduction target. Collectively, this work resolves key discrepancies in our understanding of ipRGC phototransduction pathways in multiple subtypes and adds to mounting evidence that ipRGC subtypes employ diverse phototransduction cascades to fine-tune cellular responses for downstream behaviors.

## Introduction

Light is a pervasive and important regulator of physiology and behavior across timescales that range from milliseconds to days. While rod and cone photoreceptors are primarily responsible for rapid, spatially discrete light signals, melanopsin phototransduction within the M1-M6 intrinsically photosensitive retinal ganglion cell (ipRGCs) subtypes spatially and temporally integrates environmental light signals to impact diverse behaviors over significantly longer timescales (Berson et al., 2002; Hattar et al., 2002). M1 ipRGCs project to non-image forming brain regions to influence subconscious non-image forming functions such as circadian photoentrainment, the pupillary light reflex, learning, and mood (Hattar et al., 2003; Mrosovsky and Hattar, 2003; Lucas et al., 2003; Panda et al., 2003; Altimus et al., 2008; Lupi et al., 2008; Göz, et al., 2008; Gooley et al., 2012; LeGates et al., 2012; Fernandez et al., 2018; Rupp et al., 2019; Sondereker et al., 2020; Aranda and Schmidt, 2021). M2-M6 ipRGCs primarily innervate brain areas involved in conscious visual perception and are necessary for proper contrast sensitivity (Ecker et al., 2010; Estevez et al., 2012; Zhao et al., 2014; Schmidt et al., 2014; Stabio et al., 2018; Quattrochi et al., 2019; Aranda and Schmidt 2021). Despite diverse behavioral functions, one common feature across ipRGC subtypes is that all require melanopsin (Opn4) phototransduction for their intrinsic light sensitivity. Indeed, melanopsin null mutant animals show deficits in both image-forming and non-image forming behaviors (Panda et al., 2002; Ruby et al., 2002; Lucas et al., 2003; Schmidt et al., 2014), highlighting the important role melanopsin plays across diverse cell types and behaviors.

Melanopsin shares greater sequence homology with invertebrate rhodopsins than vertebrate opsins, which led to an early expectation that melanopsin phototransduction in all ipRGC subtypes would work through an identical, invertebrate-like signaling pathway involving activation of a Gq/PLC-based cascade that opens canonical transient receptor potential (TRPC) channels such as TRPC 3, 6, or 7 (Provencio et al., 1998, 2000, 2002; Koyanagi et al., 2005; Koyanagi and Terakita, 2008; Graham et al., 2008; Hartwick et al., 2007; Perez-Leighton et al., 2011; Warren et al., 2006; Xue et al., 2011). However, though M1 ipRGCs use the predicted Gq/PLC based cascade to open only TRPC6/7 channels, phototransduction in non-M1 ipRGCs has been reported to rely less exclusively on TRPC channels and to also target additional channel types (Warren et al., 2006; Hartwick et al., 2007; Graham et al., 2008; Xue et al., 2011; Perez-Leighton et al., 2011; Sonoda et al., 2018; Jiang et al., 2018; Contreras et al., 2021). In M2 cells, for example, melanopsin phototransduction is reported to open both TRPC channels and Hyperpolarization-activated Cyclic Nucleotide-gated (HCN) channels (Perez-Leighton et al., 2011; Jiang et al., 2018; Contreras et al., 2021). In M4 cells, researchers have reached conflicting conclusions about the identity of the melanopsin transduction channel(s). Our previous findings in M4 cells point to closure of potassium leak channels by melanopsin phototransduction and a minor contribution from TRPC3/6/7 channel opening, while a concurrent study concluded that melanopsin phototransduction leads to opening of HCN channels in M4 cells with no contribution from TRPC3/6/7 channels (Sonoda et al., 2018; Jiang et al., 2018; Contreras et al., 2021). Potassium channel closure versus HCN channel opening would lead to distinct, laregely opposing impacts on M4 cell physiology and signaling, and thus a major goal of this study was to resolve the role of HCN and TRPC channels in ipRGC phototransduction in M4 cells, as well as to re-assess their role in M2 ipRGCs.

In this study, we show that HCN channels are not the M4 phototransduction channel and confirm a minor role for TRPC3/6/7 channels in bright light. Unexpectedly, we also find that HCN channels are dispensable for M2 phototransduction and identify T-type voltage-gated calcium channels as a novel, and critical, component of M2 phototransduction. Unlike M2 cells, M1 cells do not require T-type voltage-gated calcium channels for phototransduction, highlighting an important difference between the TRPC-dependent phototransduction cascades of M1 versus M2 ipRGC. Thus, M1, M2, and M4 ipRGCs each signal through distinct complements of phototransduction channels. Collectively, our findings resolve important discrepancies in our understanding of ipRGC phototransduction and add to the growing body of evidence for diverse melanopsin signaling cascades across ipRGC subtypes.

## Results

### TRPC3/6/7 channels contribute to M4 phototransduction

Two models for M4 phototransduction have been proposed. Our previous work suggests that melanopsin in M4 ipRGCs leads to potassium channel closure with a minor contribution of TRPC3/6/7 channel opening in bright light (Sonoda et al., 2018), while a separate study published concurrently proposed that HCN channels are opened by the melanopsin phototransduction pathway with no contribution from TRPC channels (Jiang et., 2018). To begin to resolve these discrepancies, we first recorded the M4 photocurrent under conditions that matched those used in Jiang et al. as closely as possible (Table 1 and see Methods). We stimulated M4 cells with brief, full-field, 50ms flashes of high photopic (6.08 × 10^15^ photons · cm^−2^ · s^−1^) 480 nm light, the highest possible intensity for our LED light source. We opted to use Infrared Differential Interference Contrast (IR-DIC) localization to initially identify the large somata of putative M4 ipRGCs because it best minimizes bleaching compared to epifluorescence. We filled each recorded cell with Neurobiotin and confirmed the identity of each M4 cell post-recording using multiple, established criteria including morphology (verified ON stratification, large somata, highly branched dendritic arbors), physiology (ON-sustained responses to increments in light), and immunolabeling for SMI-32 (Schmidt et al., 2014; Lee and Schmidt, 2018; Sonoda et al., 2018; Sonoda et al., 2020) (Figure 1A). We recorded the M4 melanopsin photocurrent in a cocktail of synaptic blockers (see Methods) at −66mV following exposure to a brief, 50ms full field flash. Control M4 cells consistently exhibited an inward photocurrent that was composed of both a relatively fast, transient component followed by a slower, larger inward current that persisted for more than 30s after termination of the 50ms light pulse (Figure 1B-1C). This small, transient component was previously noted in a subset of M2 cells but has not been observed in M4 cells in photopic light, potentially due to the previous use of lower intensity photopic stimuli (Sonoda et al., 2018) or the use of epifluorescent illumination to localize M4 cells that were subsequently recorded in high photopic light (Jiang et al., 2018, Ecker et al., 2010).

**Figure 1.**
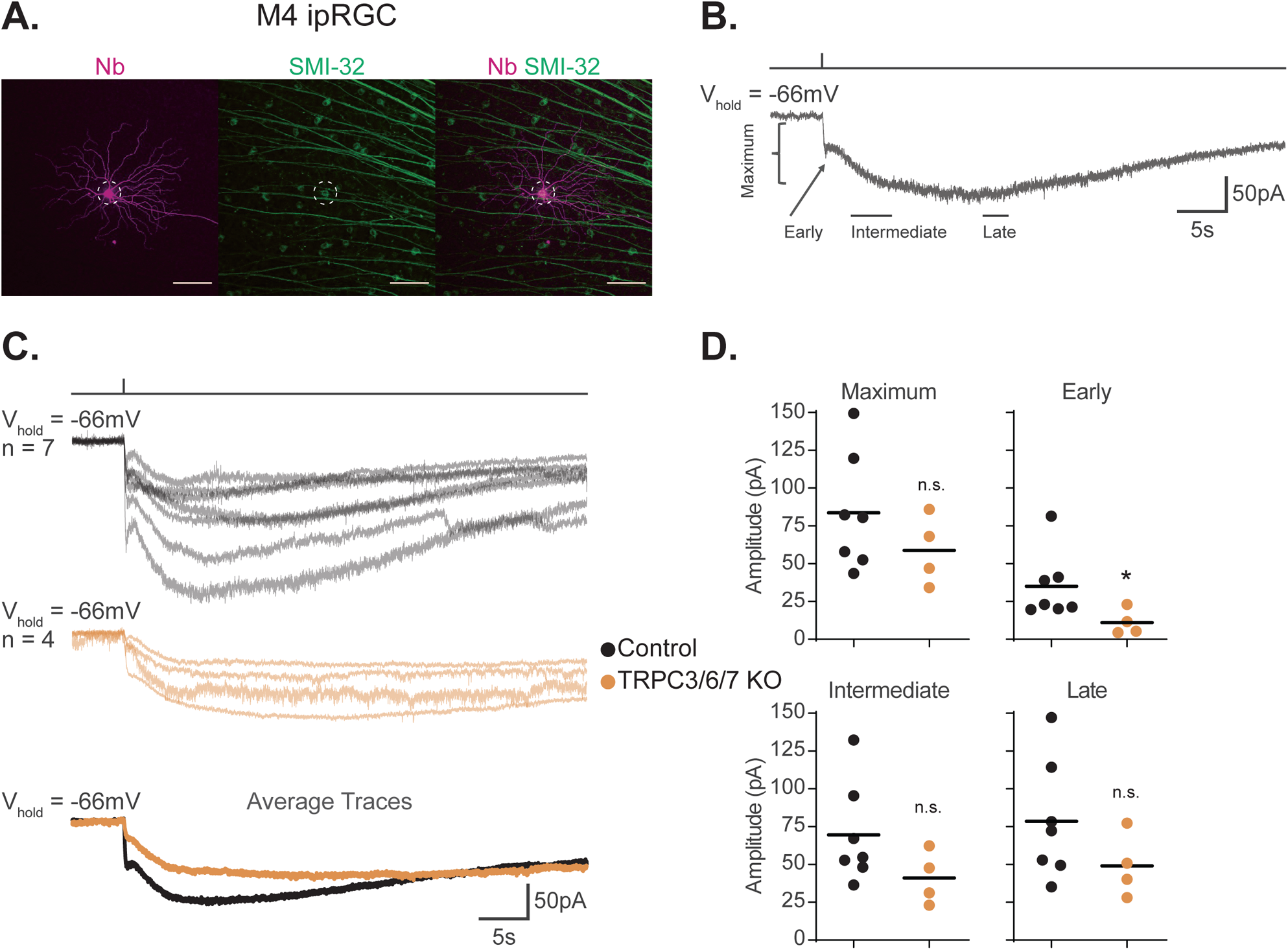
TRPC3/6/7 channels make a minor contribution to the melanopsin photocurrent in M4 ipRGCs. (A) M4 ipRGC filled with Neurobiotin (Nb, magenta) in a Control retina and immunolabeled for the M4 marker SMI-32 (green). Right panel shows merged image of overlap, M4 cell outlined with white dotted circle. Scale bar 100µm. (B) Example whole-cell voltage-clamp recording of M4 ipRGC photocurrent in a Control retina stimulated by a 50ms, full-field 480nm light (6.08 × 10^15^ photons · cm^−2^ · s^−1^) pulse in the presence of synaptic blockers. The trace is labeled with the Maximum, Early, Intermediate, and Late components used in the analysis (see methods). (C) Individual light responses of Control (top row, black, n = 7) and TRPC3/6/7 KO (middle row, orange, n=4) M4 ipRGCs. The bottom row are the overlaid averages of M4 Control (black) and TRPC3/6/7 KO (orange) light response. (D) The absolute value of the current amplitudes for light responses in panel (C) are quantified. The graphs for the Maximum, Early, Intermediate, and Late components compare the current amplitude for Control (black, n = 7) and TRPC3/6/7 KO (orange, n=4) M4 ipRGCs. The Early component of TRPC3/6/7 KO M4 ipRGCs are significantly reduced compared to Control (* p=0.0424). Recordings are in the presence of synaptic blockers. All cells were stimulated with a 50ms flash of blue (480nm) light (6.08 × 10^15^ photons · cm^−2^ · s^−1^). * p < 0.05. n.s., not significant. Analysis performed using the Mann Whitney U test (see methods)

**Table 1.** Side-by-side comparison of parameters and cell subtype identification criteria in current study and Jiang et al., 2018. Citations provided in Methods text.

|  | Current Study |  | Jiang et al., 2018 |  |
| --- | --- | --- | --- | --- |
|  | M2 | M4 | M2 | M4 |
| Mouse models | Opn4-GFP<br>Trpc3<sup>-/-</sup>; Trpc6<sup>-/-</sup>; Trpc7<sup>-/-</sup> | WT<br>Trpc3<sup>-/-</sup>; Trpc6<sup>-/-</sup>; Trpc7<sup>-/-</sup> | Opn4-tdTomato<br>Primarily: Trpc6<sup>-/-</sup>; Trpc7<sup>-/-</sup><br>*Figure 2*: Trpc1<sup>-/-</sup>; Trpc3<sup>-/-</sup>; Trpc4<sup>-/-</sup>; Trpc5<sup>-/-</sup>; Trpc6<sup>-/-</sup>; Trpc7<sup>-/-</sup> | Opn4-tdTomato<br>Primarily: Trpc6<sup>-/-</sup>; Trpc7<sup>-/-</sup><br>*Figure 2*: Trpc1<sup>-/-</sup>; Trpc3<sup>-/-</sup>; Trpc4<sup>-/-</sup>; Trpc5<sup>-/-</sup>; Trpc6<sup>-/-</sup>; Trpc7<sup>-/-</sup> |
| Cell targeting ex vivo | Epifluorescence intensity | IR-DIC/soma size and ON-sustained light response | Epifluorescence intensity/soma size | Epifluorescence intensity/soma size |
| Subtype identity | During Recording:<br>-Stratification analysis using Alexa 594<br>-Neurobiotin fill for post-hoc morphological analysis<br>Post Recording:<br>-SMI-32 negative<br>-ON-stratification<br>-Large arbors with moderate branching<br>Defining feature: M2 ipRGCs are the only ON stratifying ipRGC labeled in Opn4-GFP mice and they are SMI-32 negative. | During Recording:<br>-Stratification analysis using Alexa 594<br>-Neurobiotin fill for post-hoc morphological analysis<br>Post Recording:<br>-SMI-32 Positive<br>-ON-stratification<br>-Highly branched arbors<br>Defining feature: M4 ipRGCs are the only SMI-32 positive ipRGC, and are ON stratifying | Intracellular dye filling: Alexa 568 for morphological analysis of dendritic arbors (criteria unspecified) | Intracellular dye filling: Alexa 568 for morphological analysis of dendritic arbors (criteria unspecified) |
| Mice dark adapted | overnight |  | 3 hours |  |
| Technique | whole-cell voltage-clamp recording |  | whole-cell voltage-clamp recording |  |
| Holding potential | -66mV for all experiments unless otherwise mentioned |  | -66mV for all experiments except *Figure 4* |  |
| Internal Solution (mM) | 120 K-gluconate, 5 NaCl, 4 KCl, 10 HEPES, 2 EGTA, 4 ATP-Mg, 0.3 GTP-Na<sub>2</sub> and 7-Phosphocreatine-Tris, with the pH adjusted to 7.3 with KOH. |  | 120 K-gluconate, 5 NaCl, 4 KCl, 10 HEPES, 2 EGTA, 4 ATP-Mg, 0.3 GTP-Na<sub>2</sub> and 7-Phosphocreatine-Tris, with the pH adjusted to 7.3 with KOH. |  |
| Synaptic Blockers | 100μM DNQX, 20μM L-AP4, 100μM picrotoxin, and 20μM strychnine |  | 20μM DNQX, 50μM AP5, 100μM Hexamethonium, 100μM picrotoxin, and 1μM Strychnine |  |
| Recording Temperature | 30–32°C |  | 30–32°C |  |
| ZD7288 Conditions | Concentration: 50μM<br>Incubation time for effective HCN blockade: 5-8min<br>Incubation time driving off-target effects: 20min |  | Concentration: 50μM<br>Incubation time: not reported |  |
| Light step | 50ms |  | 200ms |  |
| Light intensity | 6.08 x 10<sup>15</sup> photons · cm<sup>-2</sup> · s<sup>-1</sup> blue LED light (480 nm) |  | White light of an intensity equivalent to 1.75 × 10<sup>18</sup> photons cm<sup>-2</sup> sec<sup>-1</sup> of 480-nm light for melanopsin (conversion done by response-matching in the linear range) |  |

We next assessed the role of TRPC3/6/7 channels in M4 phototransduction. Our previous work described a relatively minor, but detectable, role for TRPC3/6/7 channels in M4 phototransduction in photopic light (10^12^ photons · cm^−2^ · s^−1^), while others have found no contribution of these channels at higher photopic light intensities (10^18^ photons · cm^−2^ · s^−1^) (Sonoda et al., 2018, Jiang et al., 2018). To determine the role of TRPC3/6/7 channels in M4 phototransduction, we compared the intrinsic photocurrent of control M4 cells to that of M4 cells in *Trpc3^−/-^*; *Trpc6^−/-^*; *Trpc7^−/-^* (TRPC3/6/7 KO) retinas to 50ms full-field flashes of the same bright, full-field, 480 nm light (6.08 × 10^15^ photons · cm^−2^ · s^−1^) (Figure 1C-D). We analyzed the photocurrent amplitude at Early, Intermediate, and Late timepoints in the recording, as well as the maximum photocurrent amplitude (Figure 1D and see Methods). We found a significant decrease in the amplitude of the Early, transient, component in TRPC3/6/7 KO M4 cells, suggesting that this component is largely driven by TRPC3/6/7 channels. We also observed a slight, but not significant, decrease in all other quantified amplitudes. Collectively, these data are consistent with our previous findings that TRPC3/6/7 channels make a minor contribution to the M4 light response in bright light (Figure 1C-D) (Sonoda et al., 2018).

### HCN channels are not required for M4 phototransduction

We next examined the reported contribution of HCN channels to M4 phototransduction. HCN channels are a class of nonspecific cation channels opened primarily by membrane hyperpolarization whose activation voltage shifts in the presence of cyclic nucleotides (Biel et al., 2009). One compelling piece of evidence in support of melanopsin phototransduction opening HCN channels was a reduction of the M4 photocurrent following application of 50μM ZD7288, an HCN antagonist (Jiang et. al., 2018). We therefore first tested whether 50μM ZD7288 in our hands blocked both the M4 photocurrent and HCN-mediated tail current using similar recording conditions to Jiang et al., combined with rigorous post-hoc subtype identification of M4 cells. Because the incubation period for ZD7288 was not reported by Jiang et al., we first tested whether a period of 5-8 minutes incubation with ZD7288 was sufficient to abolish the HCN current in M4 ipRGCs. To do this, we recorded HCN tail currents from M4 ipRGCs first in the absence and then in the presence of 50μM ZD7288 (Figure 2A) (Chen and Yang, 2007; Van Hook and Berson, 2010). We found that treatment with 50μM ZD7288 for 5-8 minutes was sufficient to eliminate the HCN tail current and achieve full HCN channel blockade in M4 ipRGCs (Figure 2A). After confirming tail current blockade by 5-8 min of 50μM ZD7288, we then measured the photocurrent of these same M4 cells. If HCN channels are the sole target of melanopsin phototransduction in M4 cells, then this full blockade of HCN channels with 5-8 minutes of 50μM ZD7288 should eliminate the melanopsin-dependent M4 photocurrent. However, we observed no reduction in the M4 photocurrent in 50μM ZD7288 despite elimination of the HCN tail current under these conditions (Figure 2B-C). Thus, though M4 ipRGCs do express HCN channels, HCN channels are not required for melanopsin phototransduction.

**Figure 2.**
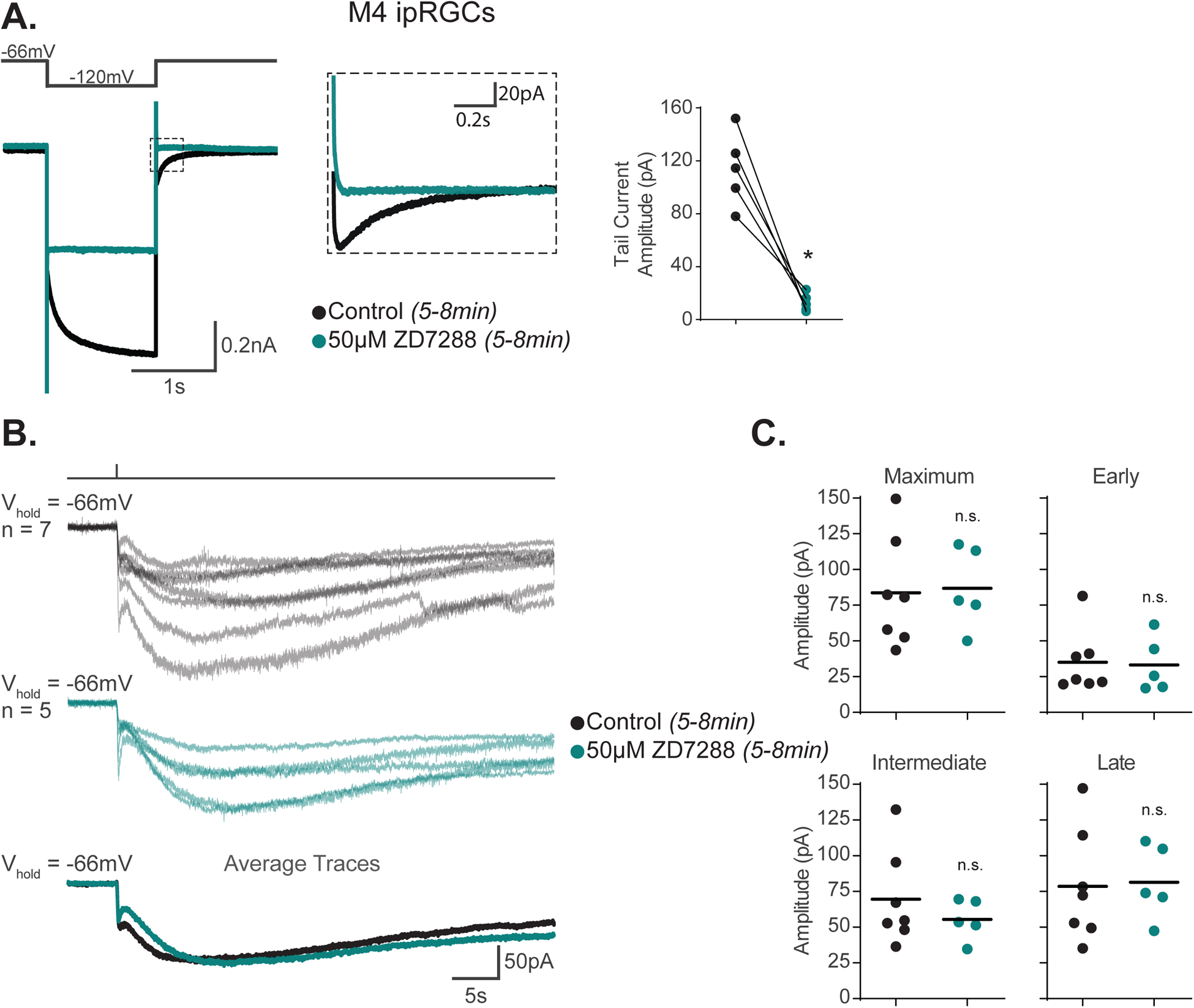
HCN channels are not required for M4 phototransduction. (A) Left, representative recording a typical Control M4 ipRGC (black), hyperpolarized from −66mV to −120mV and stepped back to the original holding potential. Cell was then incubated for 5-8min with 50μM ZD7288 (teal) and subjected to the same voltage clamp protocol. Tail currents are boxed. Middle, magnified boxed tail currents. Right, absolute value of the tail current amplitude of Control M4 cells (black, n=5) before and after application of 50μM ZD7288 for 5-8 min (teal, n=5). 50μM ZD7288 for 5-8min successfully blocked HCN mediated tail currents of M4 ipRGCs (p=0.0079). Analysis performed using Wilcoxon signed-rank test (see methods) (B) Individual light responses of M4 cells recorded in control solution (black, n=7) or M4 cells incubated with 50μM ZD7288 for 5-8min (teal, n=5, same cells for which tail current was quantified in panel A). Bottom row shows the overlaid average light response trace for Control (black) and 50μM ZD7288 for 5-8min (teal) M4 cells. (C) Maximum, Early, Intermediate, and Late absolute value amplitudes of Control (black, n=7) M4 ipRGCs or cells exposed to 5-8min of 50μM ZD7288 (teal, n=5). The photocurrent of M4 cells in 5-8min of 50μM ZD7288 is unaffected by blockade of HCN channels as shown in by the insignificant change in all the analyzed components. * p < 0.05. n.s., not significant. Analysis performed using the Mann Whitney U test (see methods)

As a second test of HCN involvement in M4 phototransduction, we compared the I-V relationship of the M4 HCN tail current versus that of the M4 photocurrent. We previously showed that the photocurrent I-V relationship of Control and TRPC3/6/7 KO M4 cells has a negative slope, reverses at the equilibrium potential for potassium channels (−90mV), and drives an increase in input resistance and cellular excitability, indicating that potassium channels are the primary target of M4 phototransduction (Sonoda et al. 2018). If melanopsin phototransduction does open HCN channels in M4 cells, then we would expect the I-V relationship of the *HCN tail current* itself to match the reported I-V relationship of the M4 *photocurrent*. However, the I-V relationship of the HCN tail current has not previously been reported for M4 ipRGCs. We therefore measured the slope and reversal potential of the M4 HCN tail current and compared it to that of the M4 photocurrent reported previously (Sonoda et al., 2018). To do this, we used a voltage clamp protocol designed to open HCN channels, and recorded the HCN tail current across multiple test potentials in the absence and then presence of 5-8 min of 50μM ZD7288 (Chen and Yang, 2007; Van Hook and Berson, 2010) (Figure 3A). We then calculated the HCN tail current amplitude by subtracting the tail current amplitude in ZD7288 from those generated in control solution at each test potential and plotted the ZD7288-sensitive current and used a linear fit to extrapolate the reversal potential (Chen and Yang, 2007; Van Hook and Berson, 2010) (Figure 3A-B). Unlike the I-V relationship of the M4 photocurrent, the M4 HCN tail current I-V relationship has a positive slope and reverses at −26 mV. While consistent with the properties of HCN channels (Biel et al, 2009), the M4 tail current I-V relationship is distinct from that of the M4 photocurrent (M4 photocurrent replotted in Figure 3B from Sonoda et al., 2018, see also Figure 7D and S6 of Sonoda et al. 2018). These distinct differences further suggest that M4 phototransduction does not open HCN channels.

**Figure 3.**
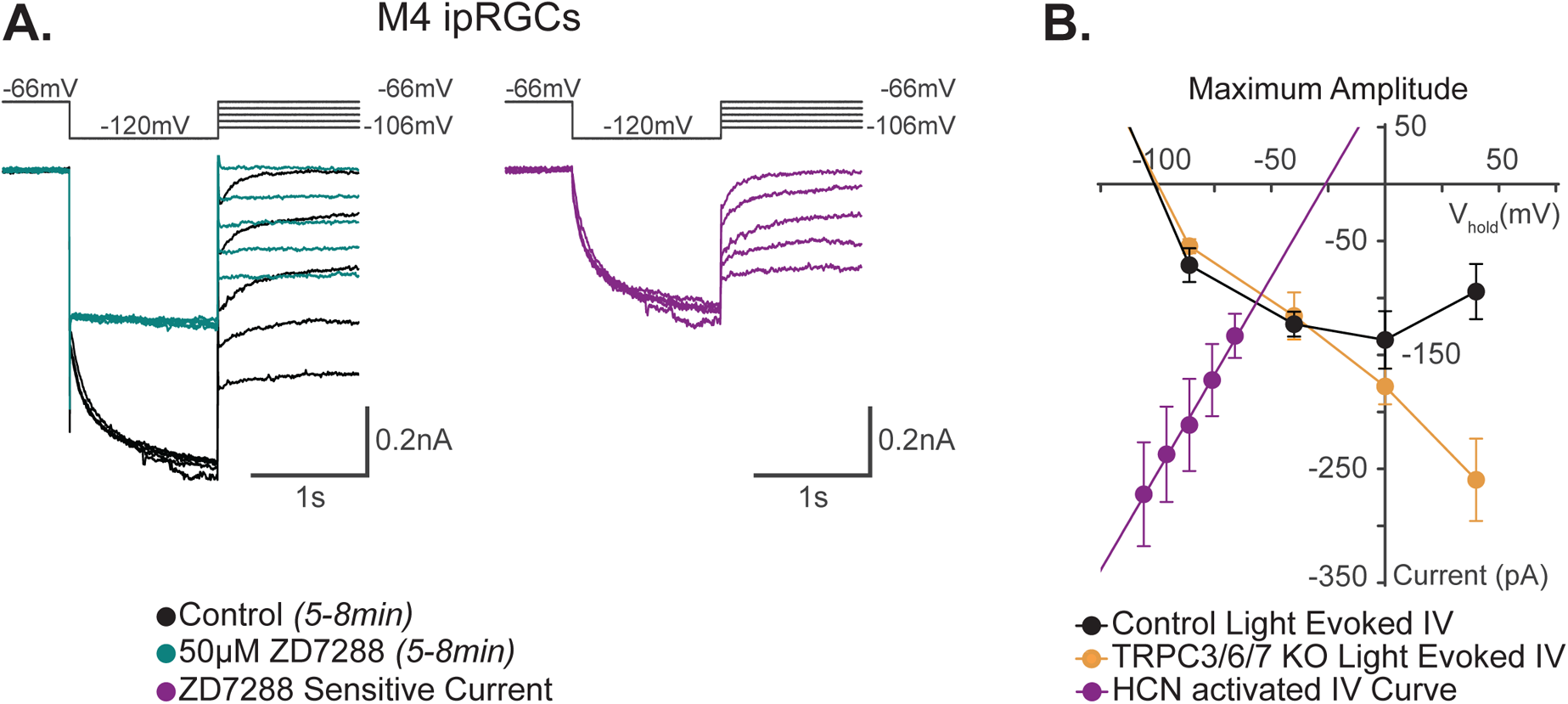
I-V relationship of M4 photocurrent is distinct from M4 HCN current. (A) Left, example Control (black) M4 cell hyperpolarized to −120mV to activate HCN channels followed by a step to various test potentials ranging from −106mV to −66mV. The voltage protocol is then repeated in the same cell following 50μM ZD7288 for 5-8min (teal). Right, the ZD sensitive currents (magenta) are obtained by subtracting the application of 50μM ZD7288 for 5-8min (teal) from Control (black). (B) Current-Voltage (I-V) relationship of M4 HCN current derived from the ZD sensitive trace (purple) in (A). A linear fit is used to extrapolate the reversal potential of HCN channels as described previously (Chen and Yang, 2007; Van Hook and Berson, 2010). The HCN mediated tail current has a positive slope and reversed at −26 mV (magenta, n=7). Data are represented as mean ± SEM. I-V relationship of HCN channels (purple) in M4 ipRGCs is compared to the 10 second light step (10^12^ photons · cm^−2^ · s^−1^) I-V relationships of the maximum photocurrent of Control (black, n=25, 5 cells/group) and TRPC3/6/7 KO M4 cells (orange, n=23 cells, 4-6 cells/group) reported by Sonoda et al., 2018. The light evoked I-V relationships for both Control (black) and TRPC3/6/7 KO (orange) M4 cells have a negative slope and reverse at −90 mV (see Figure 7D [10 second light step], and Figure S6 [100ms light step]) in Sonoda et al., 2018). This contrasts with the HCN current I-V relationship (purple) of M4 ipRGCs.

As a third test of HCN involvement in M4 phototransduction, we investigated whether light exposure increases the amplitude of the HCN tail current in M4 ipRGCs. If melanopsin phototransduction opens HCN channels, then light stimulation should result in increased HCN tail current amplitude. To test this, we compared the amplitude of the M4 HCN tail current in darkness and after 90s of bright, background light (Figure 2 – figure supplement 1; see Methods). We found no change in the M4 HCN tail current amplitude in light versus dark, suggesting that light does not modulate HCN channels in M4 cells, further arguing against HCN involvement in M4 phototransduction Figure 2 – figure supplement 1).

### Extended ZD7288 application drives off-target effects on the M4 photocurrent

Though Jiang et al. 2018 reported complete blockade of the M4 photocurrent in 50μM ZD7288, we saw no change in photocurrent amplitude in the same cells where we had confirmed this drug completely abolished HCN tail currents after 5-8 minutes incubation (Figure 2) (Jiang et al., 2018). We therefore sought to reconcile our experimental outcome with those previously described. ZD7288 has been reported to have off-target effects on other ion channels (Felix et al., 2003; Do and Bean, 2003; Sánchez-Alonso et al., 2008; Wu et al., 2012). If the reported prior blockade of the M4 photocurrent by 50μM ZD7288 was due to off-target effects, then longer incubation with 50μM ZD7288 could result in off-target effects that mimic the previously observed reduction in the M4 photocurrent. To test this, we treated M4 ipRGCs with 50μM ZD7288 for 20 minutes.

Importantly, this longer application period caused no further reduction in the HCN tail currents compared to cells incubated for 5-8 minutes, indicating that this longer incubation time does not block additional HCN channels (Figure 2 – figure supplement 2A and D). Despite identical HCN channel blockade, this longer 20 min incubation with 50μM ZD7288 now essentially abolished the M4 photocurrent and increased the input resistance (Figure 2 – supplement 2B-E). These results are consistent with potential nonspecific blockade of potassium channels by ZD7288 and provide a methodological explanation for previous conclusions of HCN involvement in M4 phototransduction (Do and Bean, 2003; Jiang et al., 2018). Collectively, our results argue against a role for HCN channels in M4 phototransduction and provide support for our previous conclusions that melanopsin phototransduction results in closure of potassium channels with a minor contribution of TRPC3/6/7 channels.

### HCN channels are not required for M2 phototransduction

Previous work reports that HCN and TRPC6/7 channels each contribute significantly to M2 phototransduction (Jiang et al., 2018). Considering the evidence against HCN involvement in M4 phototransduction, we next re-visited the role of HCN channels in M2 phototransduction. We first sought to replicate previous findings that the HCN antagonist ZD7288 partially reduces the M2 photocurrent (Jiang et al., 2018). To do this, we performed whole cell recordings of M2 ipRGCs identified under brief epifluorescent illumination in retinas of Opn4-GFP mice where only M1, M2, and M3 ipRGCs are labeled with EGFP (Schmidt et al., 2008). All cells were filled with Neurobiotin and identified as M2 ipRGCs post-recording using established criteria including: ON stratification (M2 ipRGCs are the only ON type of ipRGC labeled in Opn4-GFP mice), large dendritic arbors, and lack of SMI-32 immunolabeling (Figure 4A) (Schmidt and Kofuji, 2009; Lee and Schmidt, 2018; Lucas and Schmidt, 2019) (see methods). We first measured HCN tail currents in M2 cells before and after 5-8 minutes incubation with 50μM ZD7288 and found that the tail currents were completely abolished under these conditions, identifying an effective concentration and incubation period for full HCN channel blockade (Figure 4B). HCN tail currents of control M2 ipRGCs showed smaller, more variable HCN tail current amplitudes compared to those measured in M4 cells, consistent with previous reports (Jiang et al., 2018) (Figure 2A and 4B). We next evaluated the role of HCN channels in M2 phototransduction. If HCN channels are involved in M2 phototransduction, then application of 50μM ZD7288 should significantly reduce the M2 melanopsin photocurrent (Jiang et al., 2018). We therefore next recorded the M2 photocurrent evoked by brief, full-field, 50ms flashes of high photopic (6.08 × 10^15^ photons · cm^−2^ · s^−1^) 480 nm light in the presence or absence of 5-8 minutes of 50μM ZD7288 (Figure 4D and 4E). M2 photocurrents in control cells consistently showed a transient peak in the Early phase of the response and then a smaller, slow component that persisted for more than 30 seconds following light offset (Figure 4C and 4D). As with M4 cells, the M2 photocurrent showed no reduction following full HCN channel blockade (Figure 4D and 4E), suggesting that HCN channels are not a target of melanopsin phototransduction in M2 cells. As a second test of HCN involvement in M2 phototransduction, we tested whether light altered the tail current amplitude of M2 ipRGCs. Again like M4 cells, we found no change in the amplitude of the M2 HCN tail current recorded in background light compared to darkness (Figure 4 – figure supplement 1A), suggesting that the M2 HCN tail current is not modulated by light. This further supports the conclusion that HCN channels, though expressed on M2 cells, are not a target of M2 melanopsin phototransduction.

**Figure 4.**
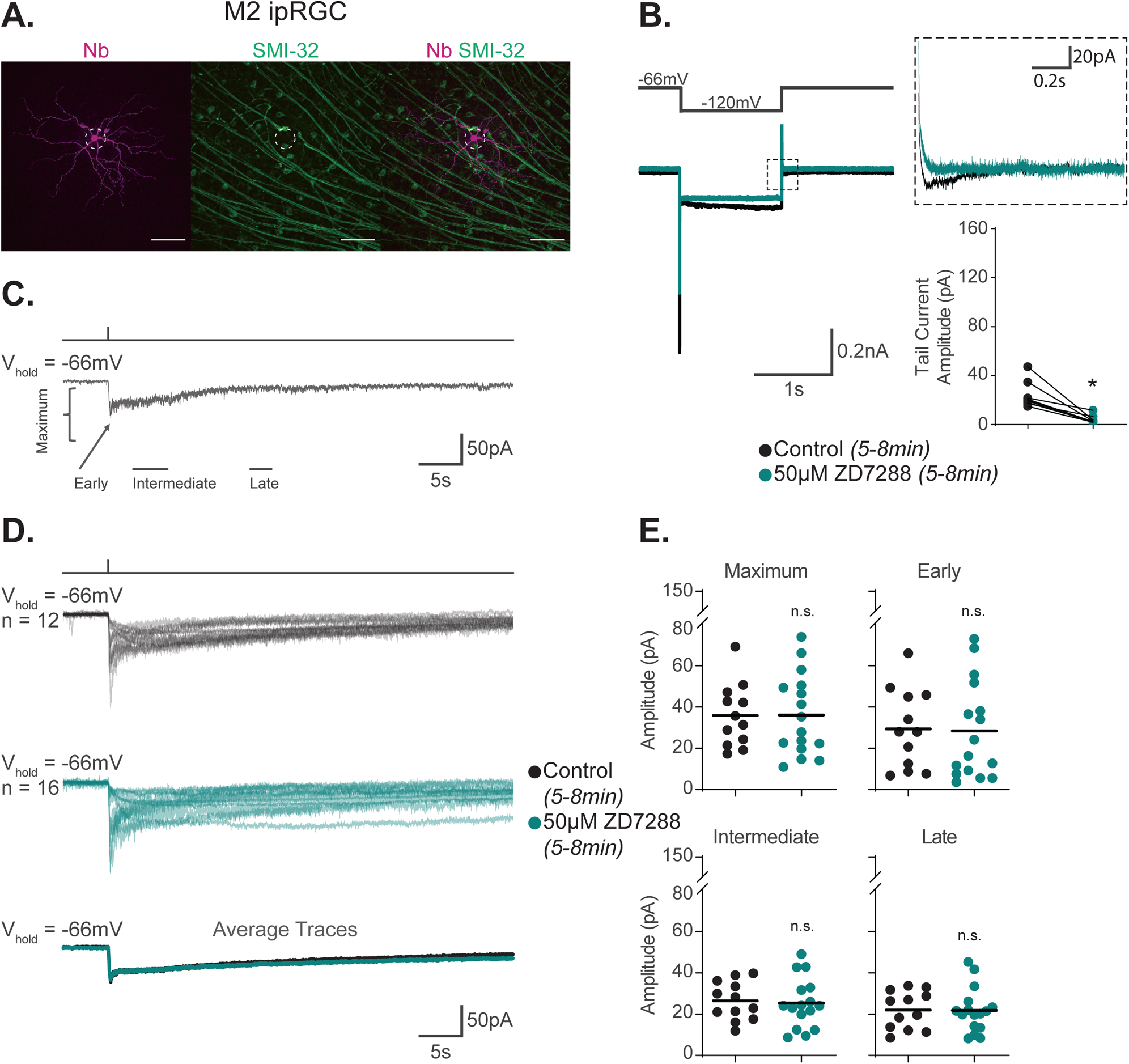
HCN channels are not required for M2 phototransduction. (A) M2 ipRGC filled with Neurobiotin (Nb, magenta) in a Control retina and immunolabeled for the M4 marker SMI-32 (green). Right panel shows merged image and lack of immunolabeling of filled cell with SMI-32, confirming identity as an M2 ipRGC. Scale bar 100μm. (B) Left, representative recording a typical Control M2 ipRGC (black), hyperpolarized from −66mV to −120mV and stepped back to the original holding potential. Cell was then incubated for 5-8min with 50μM ZD7288 (teal) and subjected to the same voltage clamp protocol. Tail currents are boxed. Middle, magnified boxed tail currents. Right, absolute value of the tail current amplitude of Control M2 cells (black, n=9) before and after application of 50μM ZD7288 for 5-8 min (teal, n=9). 50μM ZD7288 for 5-8min successfully blocked HCN mediated tail currents of M4 ipRGCs (p=0.0039). Performed statistical analysis with Wilcoxon signed-rank test (see methods). (C) Example light response of Control M2 ipRGC to a 50ms, 480nm light pulse (6.08 × 10^15^ photons · cm^−2^ · s^−1^) in the presence of synaptic blockers. Trace is labeled with the Maximum, Early, Intermediate, and Late components used in the analysis (see methods). (D) Individual light responses of Control (black, n=12) M2 cells and M2 cells incubated with 50μM ZD7288 for 5-8min (teal, n=16). Bottom row shows the overlaid average light response trace for Control (black) and 50μM ZD7288 for 5-8min (teal) M2 cells. (E) Absolute value of photocurrent amplitudes quantified for cells in (C). Photocurrent of M2 cells in 50μMZD7288 for 5-8min is unaffected despite full blockade of HCN channels shown in (B). * p < 0.05. n.s., not significant. Performed statistical analysis with Mann Whitney U test (see methods).

### Extended ZD7288 application drives off-target effects on the M2 photocurrent

Our findings suggest that HCN channels are not opened by melanopsin phototransduction in M2 ipRGCs. However, previous work did report blockade of the melanopsin photocurrent in M2 ipRGCs following incubation with ZD7288 (Jiang et al., 2018). Given our findings that ZD7288 incubation can have off-target effects that decrease the M4 photocurrent, (Figure 2 – figure supplement 2), we postulated that prolonged application to high concentrations of ZD7288 may likewise reduce the M2 photocurrent via off-target effects (Felix et al., 2003; Do and Bean, 2003; Sánchez-Alonso et al., 2008; Wu et al., 2012). To test this, we measured the photocurrent of M2 ipRGCs following 20 min incubation with 50μM ZD7288. Despite no additional reduction in HCN current, this longer incubation period resulted in a ~50% reduction in the maximum amplitude of the photocurrent in M2 cells that was consistent with the previous study (Figure 4 – figure supplement 2A-D) (Jiang et al., 2018). These findings suggest that this additional photocurrent blockade observed in this study and in previous work, was due to off-target effects on non-HCN channels in M2 cells.

### TRPC channels are a major phototransduction target in M2 ipRGCs

Given the lack of HCN involvement in M2 phototransduction, we next sought to identify the channels involved. TRPC6/7 have been reported to contribute to the M2 photocurrent (Perez-Leighton et al., 2011; Jiang et al., 2018). We therefore sought to determine the contribution of TRPC3/6/7 channels to M2 phototransduction. To do this, we recorded the M2 photocurrent in TRPC3/6/7KO; Opn4-GFP retinas. We found that the Maximum amplitude was reduced by ~75% in TRPC3/6/7 KO M2 ipRGCs, and the Early component was essentially completely abolished (Figure 5A-B). Additionally, we observed no further reduction in the small remaining photocurrent of TRPC3/6/7 KO M2 following 5-8 min incubation with the HCN antagonist ZD7288 (50μM) (Figure 5 – figure supplement 1A-C). Furthermore, the HCN tail current was not significantly increased by light exposure in TRPC3/6/7 KO M2 ipRGCs (Figure 4 – figure supplement 1). Thus, as with WT M2 phototransduction, we find no evidence of HCN involvement in TRPC3/6/7 KO M2 ipRGCs.

**Figure 5.**
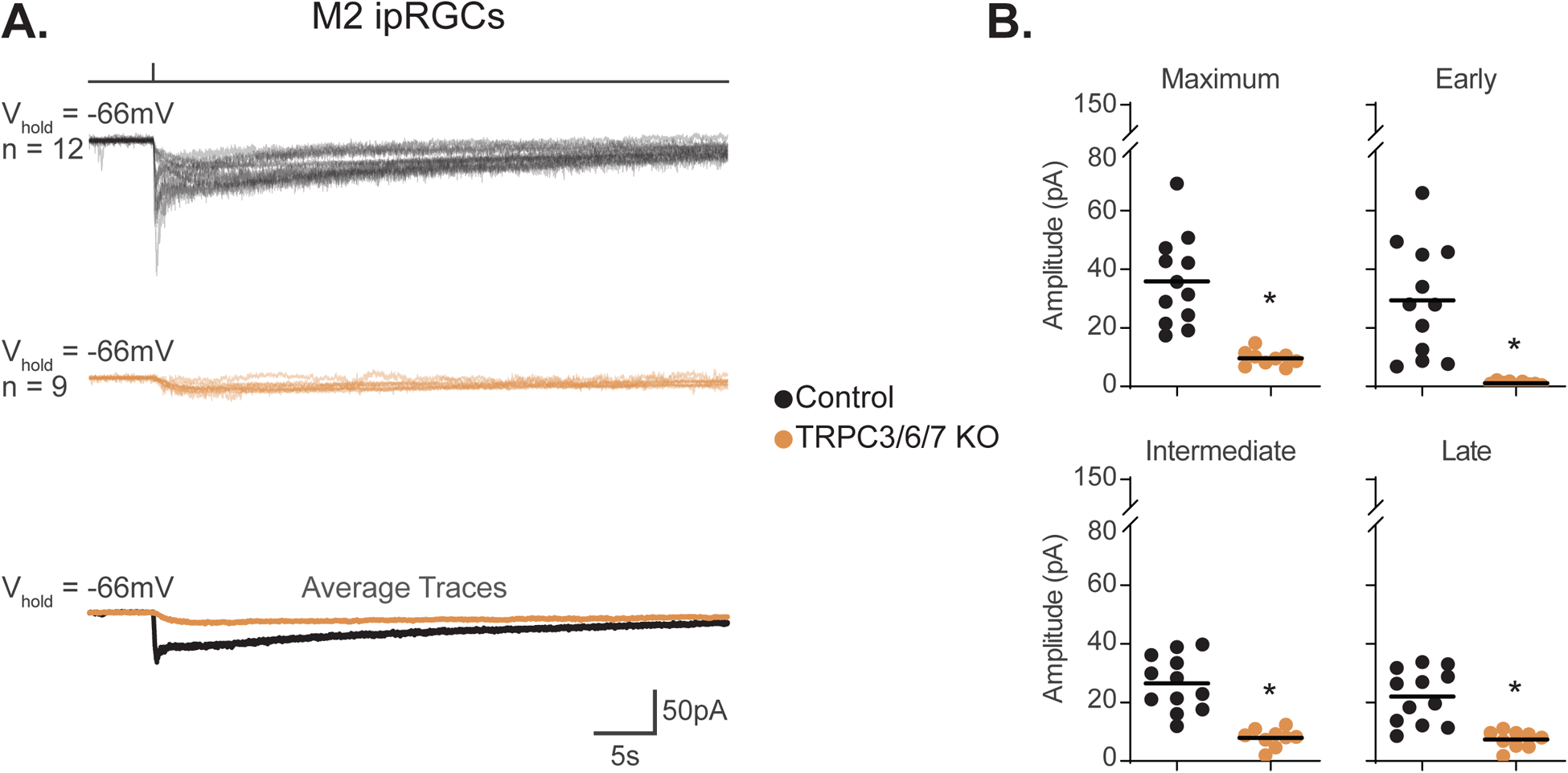
TRPC3/6/7 channels are a major phototransduction target in M2 ipRGCs. (A) Individual photocurrent recordings of Control (Opn4-GFP, black, n = 12) and TRPC3/6/7 KO (Opn4-GFP; TRPC 3/6/7 KO, orange, n=9) M2 ipRGCs to a 50ms, 480nm light pulse (6.08 × 10^15^ photons · cm^−2^ · s^−1^) in the presence of synaptic blockers. Bottom row shows the overlaid average light response trace for Control (black) and TRPC3/6/7 KO (orange). (B) Absolute value of photocurrent amplitudes quantified for cells in (A). The maximum photocurrent was significantly reduced at all phases in TRPC3/6/7 KO M2 cells (*p<0.0001). Analysis performed with Mann Whitney U test (see methods).

Our data show that TRPC channels are a major target of melanopsin phototransduction in M2 ipRGCs (Figure 5A-B), but the full I-V relationship of the M2 photocurrent has never been reported. We therefore sought to generate I-V curves for the Maximum, Early, Intermediate, and Late components. To do this, we measured the photocurrent of M2 ipRGCs at multiple holding potentials from −106mV to +34mV following stimulation with brief, full-field, 50ms flashes of high photopic (6.08 × 10^15^ photons · cm^−2^ · s^−1^) 480 nm light (Figure 6A-B, Figure 6 – figure supplement 1). In control M2 ipRGCs, the inward current was the largest at −106mV (Figure 6A and Figure 6 – figure supplement 1A) and decreased at more depolarized holding potentials (Figure 6A and Figure 6 – figure supplement 1A). The light evoked I-V curves for each component had a positive slope, consistent with an increase in conductance via channels opening (Figure 6B). All photocurrent components in control M2 ipRGCs reversed between +10mV and +25mV, suggesting contributions from cation channels that are more permeable to sodium or calcium, such as TRPC channels (Figure 6B). In TRPC3/6/7 KO retinas, I-V relationships reversed between 0 and +10mV, and the Maximum, Intermediate, and Late component amplitudes were reduced at more hyperpolarized, physiologically relevant, potentials while the Early component was reduced at all potentials (Figure 6). Notably, HCN tail currents in M2 ipRGCs reversed at −32mV, which is distinct from that of the photocurrent reversal at +19mV for control and ~0mV for TRPC3/6/7 KO M2 cells (Figure 7A-B), further arguing against HCN involvement in M2 phototransduction.

**Figure 6.**
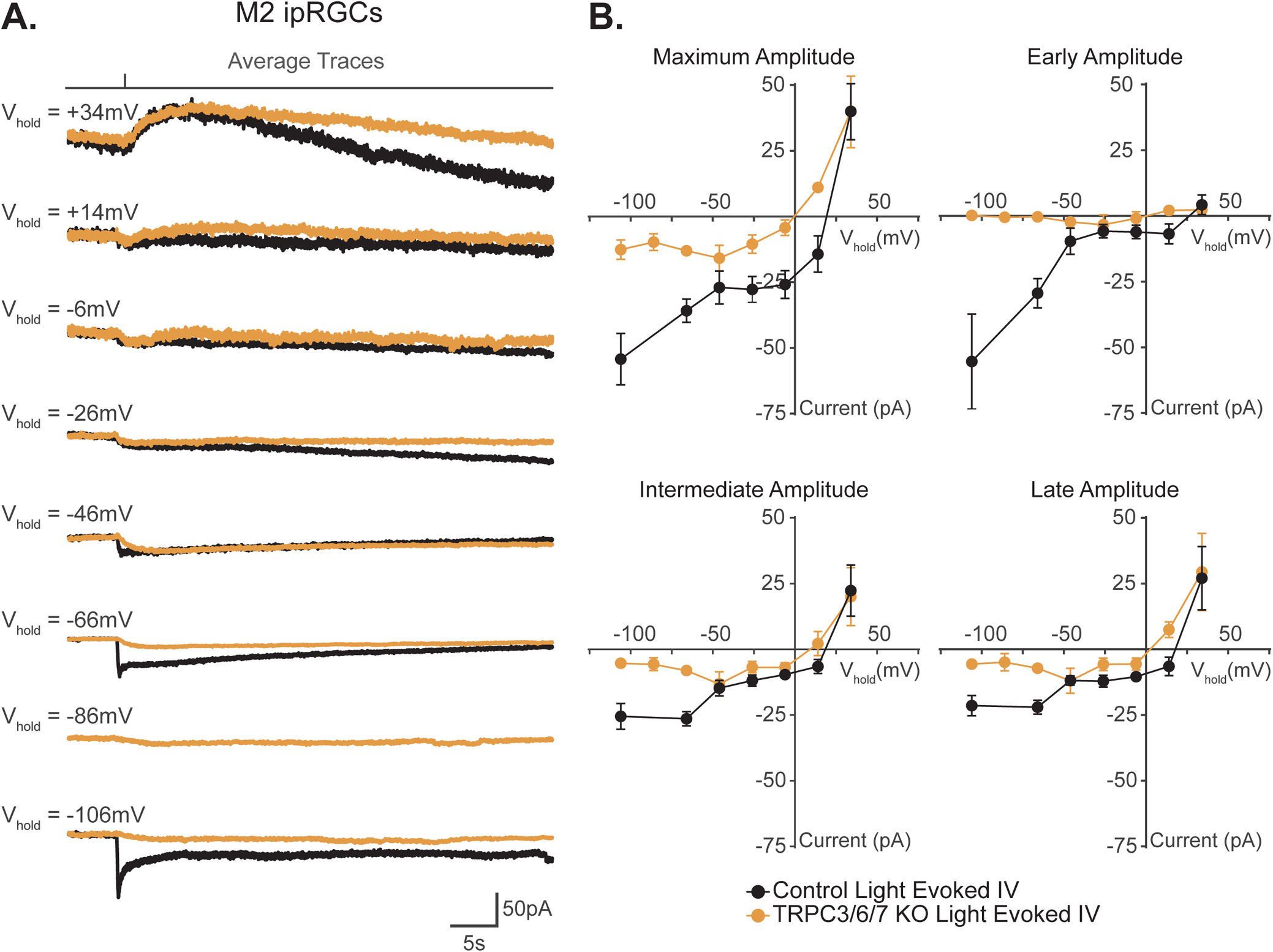
I-V relationship of M2 photocurrent. (A) Average photocurrent traces for Control (black) and TRPC3/6/7 KO (orange) M2 ipRGCs at various holding potentials (−106 mV to +34 mV) to a 50ms, 480nm light pulse (6.08 × 10^15^ photons · cm^−2^ · s^−1^) in the presence of synaptic blockers. Individual traces for all cells are shown in Figure 6 – figure supplement 1. (B) Photocurrent amplitudes for all Control (black) and TRPC3/6/7 KO (orange) M2 ipRGCs at various holding potentials for Maximum, Early, Intermediate, and Late components (black, n=52, 4-12 cells/group) and TRPC3/6/7 KO M2 ipRGCs (orange, n=43, 2-9 cells/group). I-V relationships for Control M2 cells reverse between +10-25mV and have a positive slope. Data are represented as mean ± SEM.

**Figure 7.**
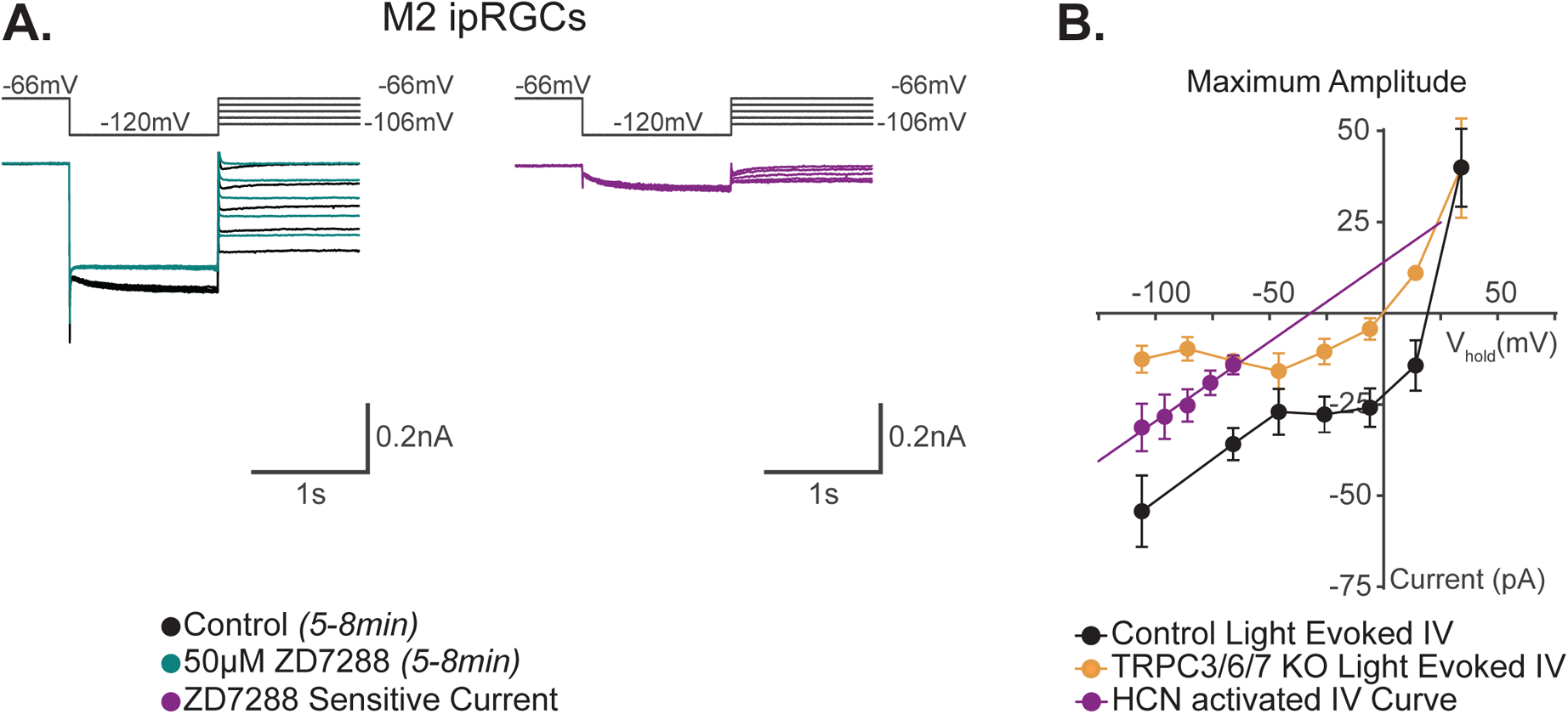
I-V relationship of M2 photocurrent is distinct from M2 HCN current. (A) Left, example Control (black) M2 cell hyperpolarized to −120mV to activate HCN channels followed by a step to various test potentials ranging from −106mV to −66mV. The voltage protocol is then repeated in the same cell following 50μM ZD7288 for 5-8min (teal). Right, the ZD sensitive currents (magenta) are obtained by subtracting the application of 50μM ZD7288 for 5-8min (teal) from Control (black). (B) I-V relationship of M2 HCN current derived from the ZD sensitive trace (purple) in (A). A linear fit is used to extrapolate the reversal potential of HCN channels. All data are represented as mean ± SEM. The HCN mediated tail current (purple, n=5) reversed at −32 mV and has a positive slope. The I-V relationship of HCN channels (purple) in M2 ipRGCs is compared to light evoked IV curves derived from the maximum component amplitudes for Control (black, n=52, 4-12 cells/group) and TRPC3/6/7 KO (orange, n=43, 2-9 cells/group) cells in Figure 6B. The reversal of HCN mediated tail current at −32mV (Figure 7) is distinct from the photocurrent reversals at +19 mV in Control M2 cells.

### Voltage-gated calcium channels are a major target of melanopsin phototransduction in M2 ipRGCs

M1 and M2 ipRGCs heavily rely on TRPC channels for phototransduction, a commonality that would predict similar photocurrent I-V relationships. However, we were surprised to see that the I-V relationship of M1 and M2 ipRGCs differed in shape and reversal potential, with M1 cells having a more linear I-V relationship that reverses at 0mV compared to a more complex I-V relationship that reverses +19 mV for M2 cells (Figure 8A) (Warren et al., 2006; Hartwick et al., 2007; Graham et al., 2008; Xue et al., 2011; Perez-Leighton et al., 2011; Sonoda et al., 2018). This observation suggests that while both M1 and M2 cells require TRPC channels for melanopsin phototransduction, another channel may be interacting with TRPC channels in M2 ipRGCs to shift the reversal of the I-V relationship. Interestingly, Voltage Gated Calcium Channels (VGCCs), like TRPC channels, can be modulated by Gq (Bloomquist et al., 1988; Scott et al., 1995; Niemeyer et al., 1996; Bertaso et al., 2003; Panda et al., 2005; Qiu et al., 2005; Hildebrand et al., 2007; Warren et al., 2006; Graham et al., 2008; Xue et al., 2011; Keum, et al., 2014; Sonoda et al., 2018; Jiang et al., 2018). Moreover, in other cell types several TRPC subunits have been shown to interact with VGCCs (Soboloff et al., 2005; Onohara et al., 2006; Yan et al., 2009; Perissinotti et al., 2021). Therefore, we hypothesized that VGCCs are involved in M2 phototransduction. To test this, we recorded the photocurrent of M2 ipRGCs in control solution and in the presence of a cocktail of VGCC antagonists (see methods) (Figure 8B and C). Application of these antagonists resulted in a significant reduction of the overall M2 photocurrent similar the that seen in TRPC3/6/7 KO M2 cells, indicating that VGCCs are required for M2 melanopsin phototransduction (Figure 8B, 8C). Importantly, we measured no change in the M1 photocurrent under blockade of VGCCs, indicating that VGCCs are not required for melanopsin phototransduction in M1 ipRGCs and their contribution may account for some of the observed differences in the M1 versus M2 photocurrent I-V relationships (Figure 8D and 8E).

**Figure 8.**
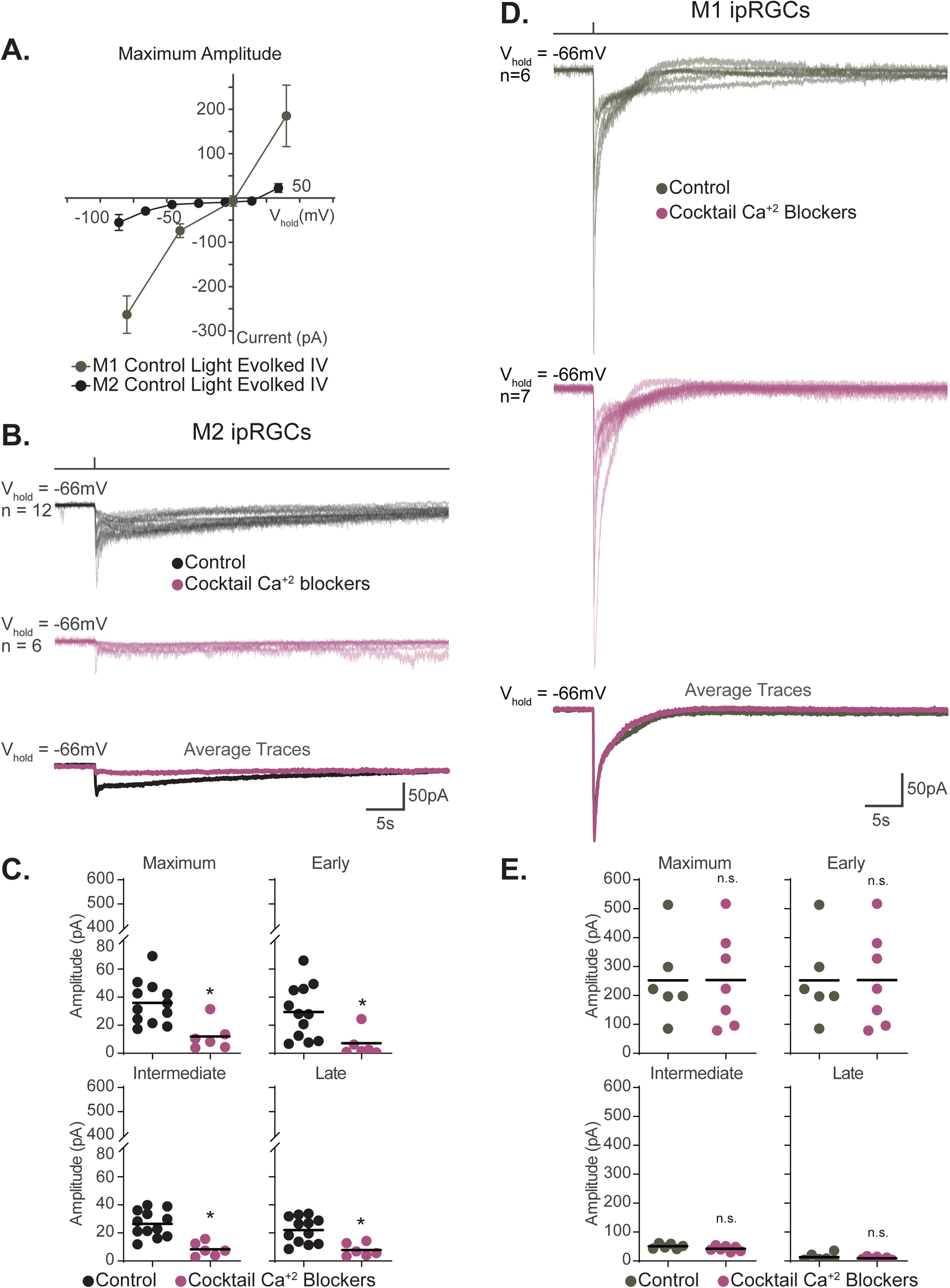
Voltage-gated calcium channels are required for M2, but not M1, phototransduction. (A) Light evoked I-V relationships for the maximum amplitude of the photocurrent in Control M2 (black, n=52, 4-12 cells/group, from Figure 6B) and M1 (grey, n = 19, 4-6 cells/group, replotted from Sonoda et al., 2018) ipRGCs. Control M2 ipRGCs (black) reversal is +19 mV compared to 0mV in Control M1 cells (grey), despite major contribution of TRPC channels to M1 and M2 phototransduction. M1 photocurrents are in response to a 10 second blue (480nm) light step (10^12^ photons · cm^−2^ · s^−1^) reported in Sonoda et al., 2018. Data are represented as mean ± SEM. (B) Photocurrent of Control (black, n=12) M2 ipRGCs in synaptic blocker cocktail and M2 ipRGCs incubated in a cocktail also containing voltage gated calcium channel antagonists (plum, n=6). Bottom row, overlaid average light response traces for Control (black) and cells in a Cocktail of Calcium Blockers (plum). VGCC antagonists were dissolved in the synaptic blocker cocktail and consisted of: 10μM nifedipine, 5μM nimodipine, 400nM ω-agatoxin IVA, 3μM ω-conotoxin GVIA, 3nM SNX-482, and 10μM mibefradil dihydrochloride. (C) Absolute value of the photocurrent of cells in (B) are quantified. The Maximum (p=0.0013), Early (p=0.0047), Intermediate (p=0.0004), and Late (p=0.0047) component amplitudes of the photocurrent are significantly reduced in the M2 ipRGCs in the presence of a Cocktail of Calcium Blockers compared to Control. (D) Photocurrent of Control (grey, n =6) M1 ipRGCs in synaptic blocker cocktail and M1 ipRGCs incubated in a cocktail also containing voltage gated calcium channel antagonists (plum, n=7). Bottom row, overlaid average light response traces for Control (black) and cells in a Cocktail of Calcium Blockers (plum). VGCC antagonists were dissolved in the synaptic blocker cocktail and consisted of: 10μM nifedipine, 5μM nimodipine, 400nM ω-agatoxin IVA, 3μM ω-conotoxin GVIA, 3nM SNX-482, and 10μM mibefradil dihydrochloride. (E) Absolute value of photocurrent for cells in (D) are quantified. Note that the Maximum and Early amplitude graphs are identical because the Early amplitude was always the maximum for M1 cells. There is no significant difference in any of the photocurrent components in M1 cells treated with a Cocktail of Calcium Blockers. All recordings for M1 and M2 ipRGCs were made in Control (Opn4-GFP) retinas in response to a 50ms flash of blue (480nm) light (6.08 × 10^15^ photons · cm^−2^ · s^−1^) and in the presence of synaptic blockers. * p < 0.05. n.s., not significant. Performed statistical analysis with Mann Whitney U test (see methods).

### T-type voltage-gated calcium channels interact with TRPC channels in M2 ipRGCs

Next, we sought to determine what type or types of VGCC is/are required for M2 phototransduction. Single cell RNA sequencing data show that T-type voltage gated channels are highly expressed in M2 ipRGCs (Tran et al., 2019). Therefore, we tested whether blockade of T-type VGCCs reduced the M2 photocurrent. We found that the T-type blocker Mibefradil Dihydrochloride (10μM) significantly reduced the M2 photocurrent, and matched the reduction seen in the full cocktail of VGCC antagonists, suggesting that T-type VGCCs are required for melanopsin phototransduction in M2 cells (Figure 9A-B). In support of this, the M2 photocurrent was unaffected under blockade of all VGCC types *except* T-type channels (Figure 9 – figure supplement 1).

**Figure 9.**
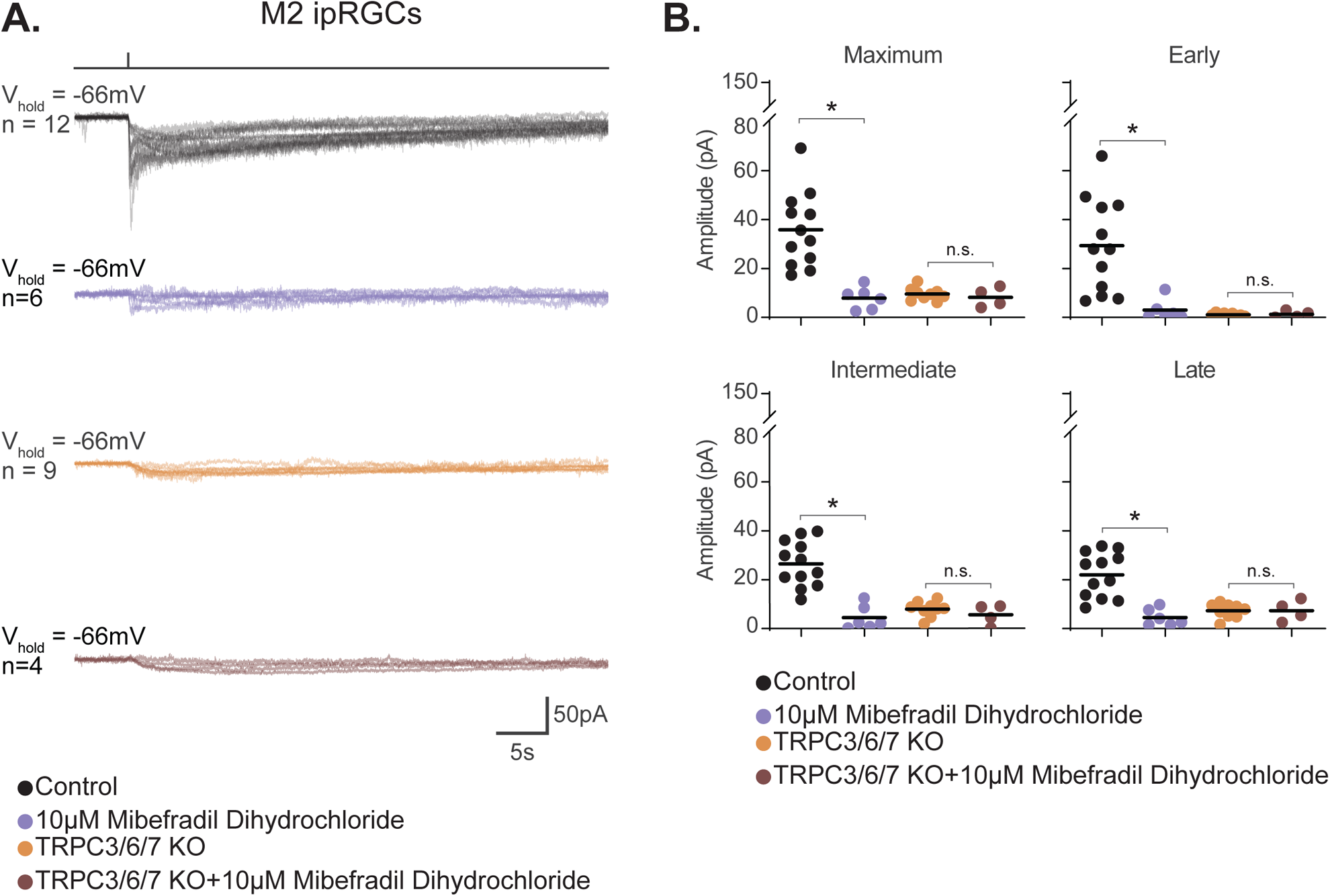
T-type voltage-gated calcium channels are required for M2 phototransduction. (A) Photocurrent of Control (black, n=12) and TRPC3/6/7 KO (orange (n=9) M2 cells recorded in synaptic blockers alone. Control and TRPC3/6/7KO M2 cells incubated in the T-type VGCC antagonist 10μM Mibefradil Dihydrochloride. Control (purple, n=6) and TRPC3/6/7 KO (brown, n=4) M2 cells in synaptic blockers plus 10μM Mibefradil Dihydrochloride. Cells were stimulated with a 50ms flash of blue (480nm) light (6.08 × 10^15^ photons · cm^−2^ · s^−1^). (B) Absolute value of photocurrent amplitudes for cells recorded in (A). The Maximum (p=0.0001), Early (p=0.0008), Intermediate (p=0.0002), and Late (p=0.0002) component amplitudes of the photocurrent are significantly reduced in the M2 ipRGCs in the presence of 10μM Mibefradil Dihydrochloride (purple, n=6). No further reduction of photocurrent was observed when TRPC3/6/7 M2 cells were incubated with 10μM Mibefradil Dihydrochloride, indicating that TRPC3/6/7 channels and T-type VGCCs are acting in the same pathway. *p<0.05. Analysis performed with Mann Whitney U test (see methods).

Intriguingly, the M2 photocurrent in TRPC3/6/7 KO M2 cells was not further reduced by Mibefradil incubation, suggesting that T-type channels and TRPC3/6/7 channels act through a common pathway (Figure 9A-B). Of note, though the degree of Mibefradil Dihydrochloride blockade mimicked that seen in TRPC3/6/7 KO M2 cells, we noted that a small transient component remained in control M2 cells under T-type blockade that is not present in TRPC3/6/7 KO M2 cells, suggesting that some current through TRPC channels may remain under T-type channel blockade in Control M2 cells.

## Discussion

Collectively, our findings identify previously unknown components of the melanopsin phototransduction cascade in M2 ipRGCs and reconcile multiple models of melanopsin phototransduction across ipRGC subtypes (Figure 10). Using rigorous, established ipRGC subtype identification criteria, we find no role for HCN channels in M2 or M4 ipRGCs. Though both subtypes express HCN channels, the melanopsin photocurrent in each subtype is insensitive to HCN blockade, has a distinct I-V relationship to that of HCN channels, and is not modulated by light.

**Figure 10.**
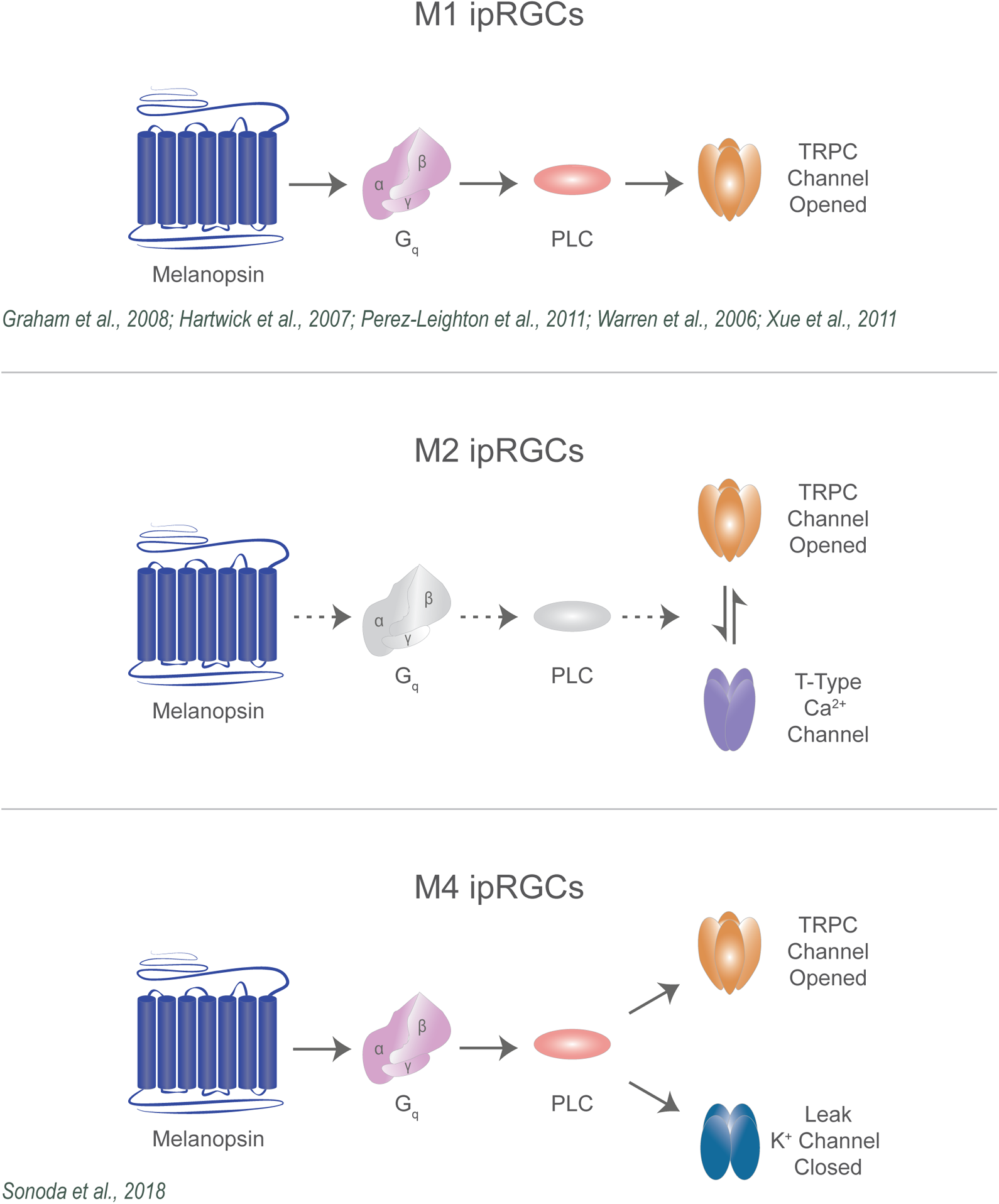
Diverse melanopsin phototransduction pathways in ipRGC subtypes. Diagram depicting melanopsin phototransduction in M1, M2, and M4 ipRGCs updated from previous work based on current findings. Figure adapted from Contreras et al., 2021. (Warren et al., 2006, Graham et al., 2008, Xue et al., 2011, Jiang et al., 2018, Sonoda et al., 2018; Perez-Leighton et al., 2011).

In our previous work, we plotted the I-V relationship of M4 ipRGCs across three experimental paradigms, using 1) whole cell patch clamp recordings to 10s light stimuli, 2) whole cell patch clamp recordings to 100ms light stimuli, or 3) nucleated patch recordings to 10s light stimuli (Sonoda et al., 2018). In all cases, the I-V relationship of the melanopsin photocurrent in WT and TRPC3/6/7 KO M4 cells had a negative slope that reversed at the potassium equilibrium potential, consistent with decreasing conductance (channel closure) through potassium channel closure via melanopsin phototransduction (Sonoda et al., 2018). Moreover, the reversal of this I-V relationship could be shifted to a newly calculated potassium equilibrium potential when the external potassium concentration was altered, further supporting a role for potassium channels in M4 phototransduction (Sonoda et al., 2018). Our previous work found evidence for a minor contribution of TRPC channels in photopic light (Sonoda et al., 2018), which we replicate in our findings here at even higher illumination. In this study, we show that the I-V relationship of the M4 *melanopsin photocurrent* is distinct from the I-V relationship of the M4 *HCN tail current*, which has the expected positive slope when HCN channels are opened, and reverses at −26 mV. Additionally, we report that the HCN antagonist ZD7288 fails to block the M4 photocurrent at a concentration and incubation period that eliminates the HCN tail current (4-8 minutes at 50μM). Additionally, light does not modulate the amplitude of the M4 HCN current. Importantly, we were able to replicate previous findings that ZD7288 blocks the M4 photocurrent, but only after a long, 20-minute incubation time. This longer incubation time did not cause additional decreases in HCN current but did increase the M4 cell input resistance, suggesting that the M4 photocurrent blockade was due to off target effects of ZD7288, which have been reported previously in other systems (Felix et al., 2003; Do and Bean, 2003; Sánchez-Alonso et al., 2008; Wu et al., 2012). Thus, while M4 ipRGCs clearly *express* HCN channels, we find no evidence that HCN channels are in fact *gated* by melanopsin phototransduction. Our findings in TRPC3/6/7 KO M4 cells also confirm that TRPC3/6/7 channels play a minor role in M4 phototransduction in bright light, which we had shown previously in lower photopic light levels (Sonoda et al., 2018), arguing against a previous report that there is no effect on M4 photocurrent amplitude in TRPC knockout M4 cells. Because much of the early transient component of the photocurrent, which depends heavily on TRPC channels, was not detected in previous work (potentially due to bleaching following epifluorescent localization), it is possible that this contribution of TRPC channels to M4 phototransduction was simply missed under highly light adapted conditions recording conditions (Jiang et al., 2018).

HCN channels had also been reported to play a role in M2 phototransduction (Jiang et al., 2018), and in light of our findings in M4 cells, we next examined the role of HCN channels in M2 phototransduction. Similar to our findings in M4 ipRGCs, 5-8min incubation period with ZD7288 effectively blocked M2 HCN channels but failed to reduce the M2 photocurrent in either Control or TRPC3/6/7 KO cells. Likewise, the HCN current reversed at a voltage distinct from the M2 photocurrent and was unaffected by light. Each of these findings argues against HCN involvement in M2 phototransduction. Only with the longer, 20 min incubation period with ZD7288 were we able to replicate the previously reported ~50% reduction in M2 ipRGCs. As with M4 cells, this blockade occurred despite no additional decrease in the HCN current, supporting the conclusion that this additional blockade was due to off-target effects on non-HCN channels (Felix et al., 2003; Do and Bean, 2003; Sánchez-Alonso et al., 2008; Wu et al., 2012). Of note, ZD7288 has been shown to block T-type calcium channels, providing further support for this interpretation (Sánchez-Alonso et al., 2008; Wu et al., 2012). Collectively, our work suggests that HCN channels are not involved in propagating melanopsin light information to downstream processing centers in the brain.

Our reevaluation of melanopsin signaling in M2 ipRGCs prompted us to look at the role of TRPC channels in M2 phototransduction. Previous studies have noted that knockout of TRPC 6 or multiple TRPC3/6/7 subunits, as well as pharmacological inhibition of TRPC channels, causes 50% or greater reduction in the M2 photocurrent suggesting that TRPC channels are a major transduction channel in M2 ipRGCs (Perez-Leighton et al., 2011; Jiang et al., 2018). In line with previous work, we found that genetic elimination of TRPC subunits 3, 6, and 7, caused a ~75% reduction in their maximum photocurrent, suggesting that TRPC channels are a major melanopsin phototransduction channel M2 cells.

Intriguingly, we observed that though both M1 and M2 ipRGCs require TRPC channels for the majority (M2) or entirety (M1) of their photocurrent, the I-V curves of M1 and M2 cells differed substantially, prompting us to probe whether other unidentified transduction channels may be involved in one or both of these phototransduction pathways. We found that blockade of T-type VGCCs resulted in a ~75% reduction in M2 photocurrent but had no effect on the M1 photocurrent, suggesting that T-type VGCCs are necessary for M2, but not M1, phototransduction. Importantly, this lack of blockade in M1 ipRGCs argues against any nonspecific effects of any VGCC blockers on TRPC channels. The similar degree of M2 photocurrent reduction by the T-type antagonist Mibefradil to that seen in TRPC3/6/7 KO M2 cells, combined with lack of TRPC3/6/7 KO M2 cell photocurrent reduction in Mibefradil suggests that TRPC channels and T-type VGCCs act through a common phototransduction pathway. The mechanisms through which this interaction occurs are unknown.

Previous studies have shown that TRPC channels and Voltage Gated Calcium Channels (VGCCs) can both be modulated by Gq signaling (Bloomquist et al., 1988; Scott et al., 1995; Niemeyer et al., 1996; Bertaso et al., 2003; Panda et al., 2005; Qiu et al., 2005; Hildebrand et al., 2007; Warren et al., 2006; Graham et al., 2008; Xue et al., 2011; Keum, et al., 2014; Sonoda et al., 2018; Jiang et al., 2018). Indeed, our findings hint that VGCCs may be downstream of, or concurrent with, TRPC channels because the small, transient component in the Early phase of the M2 photocurrent remains in Control cells recorded in Mibefradil or a cocktail of VGCC blockers. Additionally, in cell culture systems TRPC channels have been shown to modulate both L-type and T-type VGCCs (Soboloff et al., 2005; Onohara et al., 2006; Yan et al., 2009; Perissinotti et al., 2021). VGCCs have been shown to form complexes with several TRPC subunits (Perissinotti et al., 2021). TRPC3 and TRPC6 have been shown to modulate the activation of L-type channels in smooth muscle cells, and TRPC5 subunits can directly modulate T-type channels (Soboloff et al., 2005; Onohara et al., 2006; Yan et al., 2009; Perissinotti et al., 2021). Molecular work demonstrates that T-type calcium channels form complexes with TRPC1 and TRPC5 subunits (Perissinotti et al., 2021). Furthermore, TRPC1/4/5 subunits can form functional hetero-tetrameric channels with TRPC3 and TRPC6 (Strübing et al., 2003; Liu et al., 2005; Poteser et al., 2006). Intriguingly, these other TRPC subunits are all detected in M2 cells, making it possible that they form heteromeric complexes with TRPC 3/6/7 subunits that could interact with T-type VGCCs as described in other systems (Tran et al., 2019; Strübing et al., 2003; Liu et al., 2005; Poteser et al., 2006; Soboloff et al., 2005; Onohara et al., 2006; Yan et al., 2009; Perissinotti et al., 2021). The mechanisms for this remain murky in multiple systems, and more work is undoubtedly necessary to uncover TRPC and T-type VGCCs interact within the M2 melanopsin phototransduction cascade.

Our work adds to a growing literature describing the interaction between T-type calcium channels and TRPC channels, and supports divergent, TRPC-dependent phototransduction mechanisms in M1 and M2 ipRGCs. Additionally, this work resolves a key discrepancy in our understanding of M4 phototransduction and supports an emerging framework for melanopsin signaling that suggests that melanopsin phototransduction is tuned to optimize how ipRGC subtypes signal to influence their associated light-driven behaviors (Figure 10).

## Materials and Methods

### Contact for Reagent and Resource Sharing

Requests for reagents and resources should be directed to the Lead Contact, Tiffany Schmidt.

### Animals

All procedures were approved by the Animal Care and Use Committee at Northwestern University.

Both male and female mice were used with a mixed B6/129 background. All mice were between 30 and 90 days of age. For M4 cell recordings, we used WT and Trpc3-/- (Hartmann et al., 2008 RRID: MGI:3810154); Trpc6-/- (Dietrich et al., 2005 RRID: MGI:3623137); Trpc7-/- (Perez-Leighton et al., 2011 RRID: MGI:5296035) mice. For M2 and M1 cell recordings, we used Opn4-GFP (Schmidt et al., 2008) and Opn4-GFP; Trpc3-/-; Trpc6-/-; Trpc7-/- mice.

### Ex vivo Retina Preparation for Electrophysiology

All mice were dark adapted overnight and euthanized by CO2 asphyxiation followed by cervical dislocation. Eyes were enucleated, and retinas were dissected under dim red light in carbogenated (95% O2-5% CO2) Ames’ medium (Sigma-Aldrich). Retinas were then sliced in half and incubated in carbogenated Ames’ medium at 26°C for at least 30 min. Retinas were then mounted on a glass bottom recording chamber and anchored using a platinum ring with nylon mesh (Warner Instruments). The retina was maintained at 30-32°C and perfused with carbogenated Ames’ medium at a 2-4 mL/min flow.

### Solutions for Electrophysiology

All recordings were made in Ames’ medium with 23 mM sodium bicarbonate. Synaptic transmission was blocked with 100μM DNQX (Tocris), 20μM L-AP4 (Tocris), 100μM picrotoxin (Sigma-Aldrich), and 20μM strychnine (Sigma-Aldrich) in Ames’ medium. 500nM tetrodotoxin (TTX) citrate (Tocris) was added to the synaptic blocker solution for voltage-clamp experiments. For whole-cell recordings the internal solution (Jiang et al., 2018) used contained (in mM): 120 K-gluconate, 5 NaCl, 4 KCl, 10 HEPES, 2 EGTA, 4 ATP-Mg, 0.3 GTP-Na_2_ and 7-Phosphocreatine-Tris, with the pH adjusted to 7.3 with KOH. The internal solution was passed through a sterile filter with a 0.22um pore size (Sigma-Aldrich). Prior to recording, 0.3% Neurobiotin (Vector Laboratories) and 10μM Alexa Fluor 594 (ThermoFisher) were added to internal solution. The HCN antagonist, ZD7288 (Tocris) was dissolved in distilled water and added to the synaptic blockers for a final concentration of 50μM. 50μM ZD7288 was bath applied for 5-8 minutes (an incubation period we identified to fully block the HCN tail current, Figure 2A) with minimum off-target effects, or for 20 minutes, which led to additional non-specific effects on the photocurrent of M2 and M4 ipRGCs (Figure 2 – figure supplement 2 and Figure 4 – figure supplement 2). The cocktail of calcium (VGCC) blockers was added to the synaptic blockers and consisted of L-type blockers: 10μM nifedipine and 5uM nimodipine, P/Q-type blocker: 400nM ω-agatoxin IVA, N-type blocker: 3μM ω-conotoxin GVIA, R-Type blocker: 3nM SNX-482, and T-type blocker: 10μM mibefradil dihydrochloride. Nifedipine (Tocris) and nimodipine (Tocris) were dissolved in DMSO and diluted in synaptic blockers. Mibefradil dihydrochloride (Tocris), SNX-482 (Tocris), ω-conotoxin GVIA (Tocris), and ω-agatoxin IVA (Tocris) were reconstituted in water and dilute in synaptic blockers to reach the final concentration. VGCC blockers were applied for 5 minutes to minimize off-target effects.

### Light Stimulus

The blue LED light (~480 nm) was used to deliver light stimuli to the retina through a 60X water-immersion objective. The photon flux was attenuated using neutral density filters (Thor Labs). Before recording, retinas were dark-adapted for at least 5 min. The photocurrent from ipRGCs was recorded following a 50ms full field flash of bright light with an intensity of 6.08 × 10^15^ photons · cm^−2^ · s^−1^.

### Electrophysiology

The ganglion cell layer of retina was visualized using infrared differential interference contrast (IR-DIC) optics at 940nm. M4 ipRGCs, synonymous with ON-sustained alpha RGCs (Schmidt et al., 2014), were identified in IR-DIC as cells with large somata (>20 μm) and characteristic ON-sustained responses to increments in light, as described in Sonoda et al., 2018. After all cellular recordings, the identity of M4 ipRGCs was confirmed by verifying the dendrites stratified only in the ON-sublamina of the Inner Plexiform Layer (IPL) and immunolabeled with SMI-32, an M4 ipRGC marker (Schmidt et al., 2014; Sonoda et al., 2018). M1 and M2 ipRGCs were targeted in the Opn4-GFP line (M1, M2, and M3 cells labeled, M4, M5, and M6 not labeled), based on their somatic GFP signals visualized under brief epifluorescent illumination. Following recording, the identity of M1 and M2 ipRGCs were confirmed by examining dendritic stratification (M1: OFF, M2: ON) in the IPL, achieved through intracellular dye (Alexa 594) immediately following recording and confirmed post-recording by Neurobiotin fill. M2 cell dendrites stratified only in the ON-sublamina of the IPL while M1 cell dendrites stratified in the OFF-sublamina of the IPL (described in Schmidt et al., 2008 and Schmidt and Kofuji, 2009). Both M1 and M2 were confirmed negative for SMI-32 immunolabeling as described in Lee and Schmidt, 2018. For all experiments, 1 cell was recorded from each piece of retina to minimize light adaptation.

Whole cell recordings were performed using a Multiclamp 700B amplifier (Molecular devices) and fire-polished borosilicate pipettes (Sutter Instruments, 3-5 MΩ for M4 cells, 5-8 MΩ for M2 and M1 cells). All voltage traces were sampled at 10 kHz, low-pass filtered at 2 kHz and acquired using a Digidata 1550B and pClamp 10 software (Molecular devices). All reported voltages are corrected for a −13mV liquid junction potential calculated using Liquid Junction Potential Calculator in pClamp.

### Immunohistochemistry

After recording, retina pieces were fixed in 4% paraformaldehyde (Electron Microscopy Sciences) in 1X PBS overnight at 4°C. Retinas were then washed with 1X PBS for 3×30 min at room temperature (RT) and then block overnight at 4°C in blocking solution (2% goat serum in 0.3% Triton PBS). Retinas were then placed in primary antibody solution containing mouse anti-SMI-32 (1:1000, BioLegend, Cat#801701, RRID: AB_509997) in blocking solution for 2-4 days at 4°C. Retinas were washed in 1X PBS for 3×30 min at RT and transferred to secondary antibody solution containing Alexa 488 goat anti-mouse (1:1000, Thermo, Cat# A-21131, RRID:AB_2535771) and Streptavidin conjugated with Alexa 546 (1:1000, Thermo, Cat# S-11225, RRID:AB_2532130) in blocking solution overnight at 4°C. Retinas were then washed in 1X PBS for 3×30 minutes at RT and mounted using Fluoromount aqueous mounting medium (Sigma).

All images were captured using a confocal laser scanning microscope (Leica DM5500 SPE, Leica microsystems) with a 20x objective. To include whole dendrites of ipRGCs, tiled image stacks spanning the ganglion cell layer to inner nuclear layer were collected. Images were processed using Fiji (Schindelin, et al., 2012).

### Data Quantification and Analysis

All data were analyzed using custom scripts written in MATLAB (MathWorks; RRID: SCR_001622).

For voltage clamp experiments measuring the intrinsic melanopsin response to a 50ms light stimulus (6.08 × 10^15^ photons · cm^−2^ · s^−1^), ipRGCs were voltage clamped at −66mV, as in Jiang et al., 2018. Only a single cell was recorded per retina piece to ensure that the cell was not light adapted. We measured the maximum amplitude as well as the amplitude at 3 timepoints selected to represent different phases of the light response (highlighted in Figure 1B) of the photocurrent. We defined the Maximum amplitude as the current value with the greatest change from the baseline during the recording period. The Early, Intermediate, and Late time points represent the average change from baseline within the following timeframes after light onset: Early (141.7-440.4 ms), Intermediate: (2857.7-6598.2 ms), and Late (9062.3-14062.3 ms). Values were reported as the absolute value of the current (pA), for each component.

The light evoked I-V relationship for M2 cells was generated by recording the intrinsic melanopsin response for Control and TRPC3/6/7 KO M2 cells in response to a 50ms light pulse. Cells were voltage clamped at potentials from −106 to +34mV. Cells were allowed to stabilize at a given holding potentials for 1 to 3 seconds prior to light onset. Only a single cell was recorded per preparation to ensure dark adaptation state was as uniform as possible across cells/preparations. We plotted the I-V relationships for Maximum, Early, Intermediate, and Late amplitudes. The reversal potential was identified as where the I-V curve intersected the X-axis (i.e. I=0).

For experiments measuring the HCN-mediated tail current, channel activation was evoked by hyperpolarizing the cell from −66mV to −120mV (Chen and Yang, 2007; Van Hook and Berson, 2010). Tail current amplitudes were defined as the maximum change from baseline upon return to −66mV and were plotted as the absolute value of the current. For experiments assessing blockade of HCN channels, cells were exposed to 50μM ZD7288 and then subjected again to the same protocol. Blockade of HCN channels in ipRGCs was observed when the tail current amplitude was abolished (reviewed in Biel et al, 2009). Blockade of HCN channels, defined loss of tail currents, occurred between 5 and 8 minutes (5-8min) of exposure to 50μM ZD7288 for each individual cell. For experiments in Figure 2A and Figure 4B, the HCN tail current was first measured (defined as control), and then the same cell was exposed to 50μM ZD7288 for 5-8min to block HCN channels.

However, for Figure 2B and Figure 4D the photocurrent measurements were obtained from different cells: control and cells exposed to 50μM ZD7288 for 5-8 minutes. The photocurrent was measured in different cells to avoid melanopsin photobleaching. To measure the I-V relationship of HCN channels in M2 and M4 ipRGCs (as described in Chen and Yang, 2007; Van Hook and Berson, 2010), cells were hyperpolarizing −66mV to −120mV to activate the majority of HCN channels. Cells were then depolarized to various test potentials from −106mV to −66mV. This protocol was repeated in the same cells following 5-8min application of 50μM ZD7288 to ensure blockade of HCN channels. The ZD7288 sensitive tail current was derived by subtracting 5-8min ZD7288 recordings from Control. Tail current amplitudes were measured at each test potential by fitting with a single exponential. The tail current amplitudes during deactivation were plotted against each test potential to construct the I-V relationship. A linear fit was used to extrapolate the reversal potential of HCN channels. For experiments assessing light modulation of HCN channels, the tail current was first measured in the dark. The same cell was then exposed to light. The cellular membrane potential stabilized after 90 seconds of light exposure. The tail current amplitude was measured following the membrane potential stabilization in background light. For experiments assessing the effects due to prolong application of 50μM ZD7288, cells were exposed to 20 minutes of ZD7288 in whole-cell patch clamp configuration and recordings were performed following this period. Recordings for Control cells were performed 20 minutes after whole-cell was achieved to account for effects due to long recording time. Recordings with and without ZD7288 occurred in separate cells to avoid the confound of multiple light stimulations. Thus, in Figure 2 – figure supplement 2A, the tail current amplitudes (pA) of Control (20min) and 50μM ZD72288 (20min) were from different M4 cells and in Figure 4 – figure supplement 2A the photocurrent of Control (20min) and 50μM ZD72288 (20min) were from different M2 cells. Input resistance was calculated as the slope of a linear fit of the steady-state voltage deflection evoked by a series of hyperpolarizing current injections from −300 to 0pA and depolarizing current injections from 0 to 200pA.

Graphing and statistical analysis was performed using GraphPad Prism 9 software (RRID:SCR_002798). For unpaired statistical comparisons, we used a non-parametric, two-tailed Mann-Whitney U test with a Bonferroni correction. For paired statistical comparisons, we used a non-parametric, two-tailed Wilcoxon matched-pairs signed rank test. Significance was concluded when p < 0.05.

## Figure Legends

**Figure 2 – figure supplement 1.**
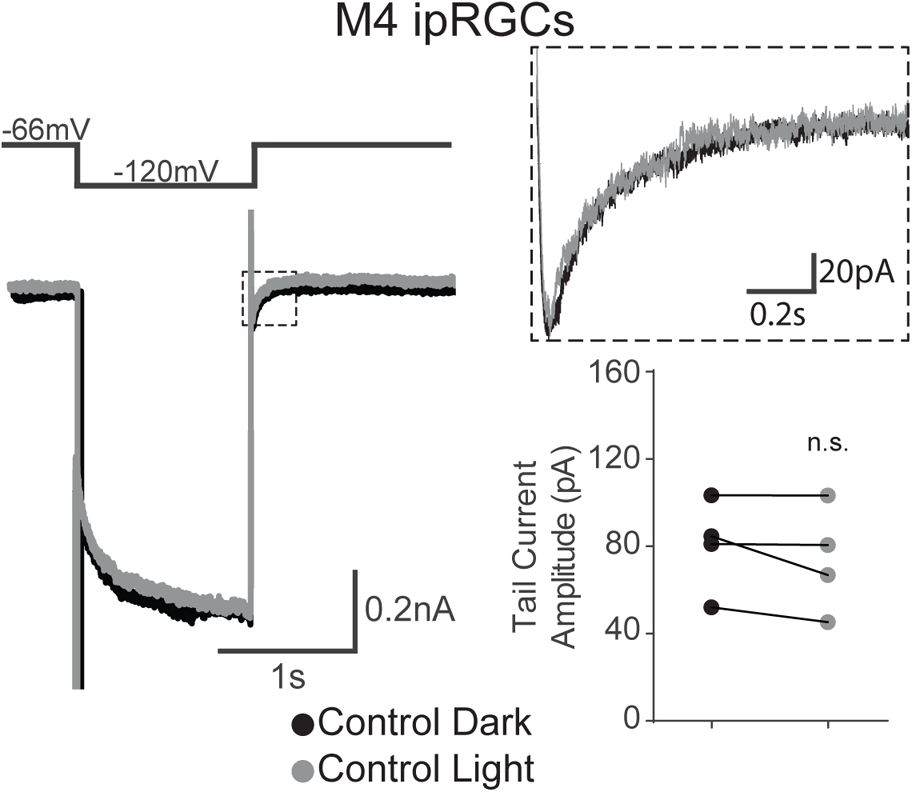
The M4 HCN tail current is not modulated by light. WT M4 ipRGC in dark (black) hyperpolarized from −66mV to −120mV and stepped back to the original holding potential. Cell was then placed in 90s of 480 nm light (6.08 × 10^15^ photons · cm^−2^ · s^−1^) (light grey) and subjected to the same voltage clamp protocol. Tail currents are boxed and expanded in inset. Absolute value of the tail current amplitude is plotted for M4 cells in dark (black, n=4) and in light (gray, n=4). There is no significant change in the tail current amplitude in light. Recordings made in presence of synaptic blockers. n.s., not significant. Performed statistical analysis with Wilcoxon signed-rank test (see methods).

**Figure 2 – figure supplement 2.**
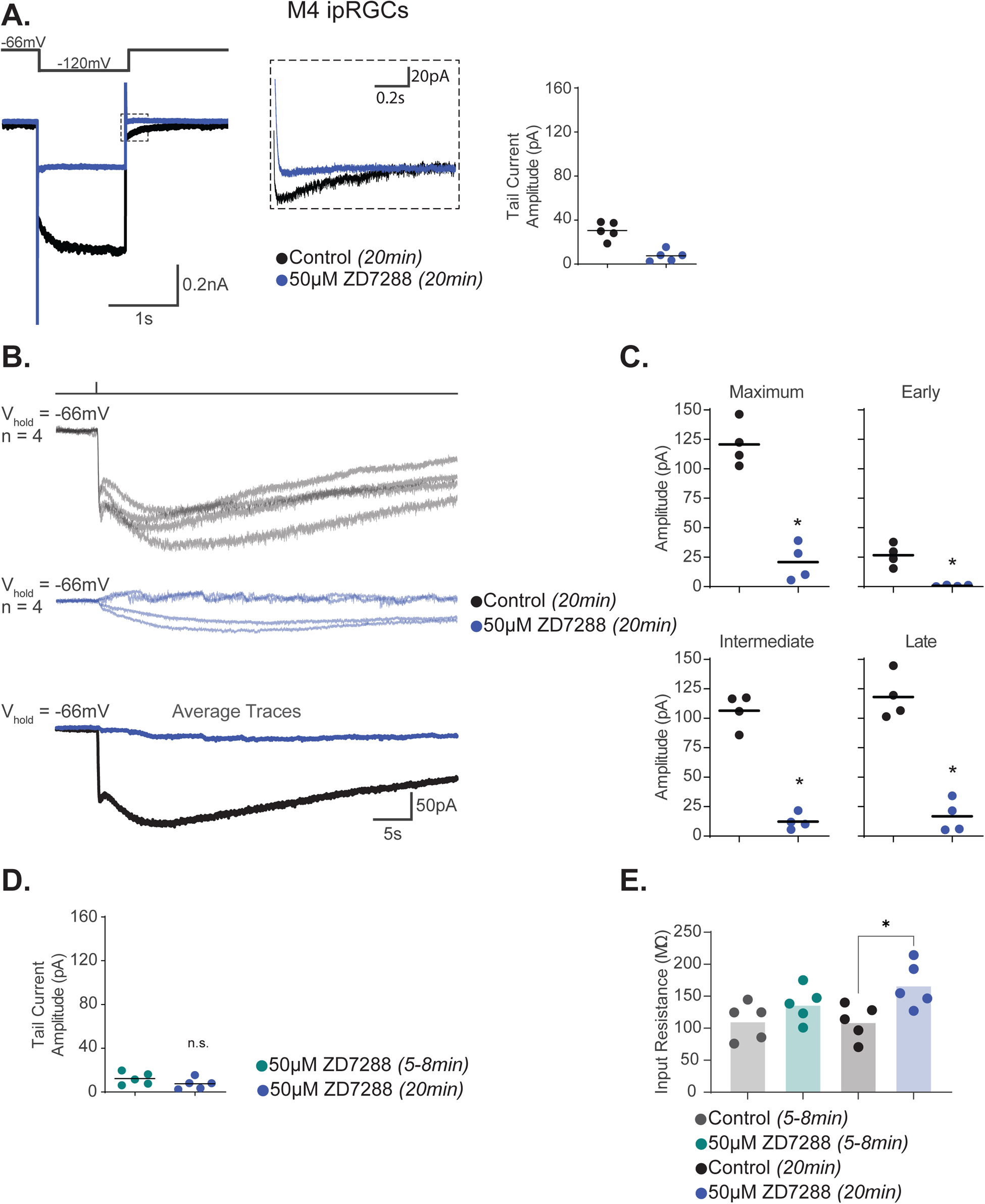
Prolonged application of ZD7288 reduces the M4 photocurrent via off-target effects. (A) WT M4 ipRGC hyperpolarized from −66mV to −120mV and stepped back to the original holding potential after 20 min in control solution (black, n=5) or after 20 min incubation 50μM ZD7288 (blue, n=5). Tail currents are boxed and expanded in inset. Absolute value of the tail current amplitude is plotted for each cell. (B) M4 photocurrent recorded after 20 min in control solution (black, n=4) or after 20 min incubation with 50μM ZD7288 (blue, n=4). Bottom row, overlaid average light response trace for each group. (C) The absolute value of the Maximum, Early, Intermediate, and Late photocurrent amplitudes of M4 cells in 20min Control solution and 20min of 50μM ZD7288 (blue, n=4). The Maximum (p=0.0286), Early (p=0.0286), Intermediate (p=0.0286), and Late (p=0.0286) photocurrent amplitudes of M4 cells in 20min of 50μM ZD7288 are significantly reduced. This recapitulates the results noted in Jiang et al., 2018. (D) Absolute value of HCN tail currents are similar after 5-8 vs. 20 min. incubation with 50μM ZD7288, indicating no further blockade of HCN channels with longer incubation. (E) Input resistance of M4 cells after 5min in Control solution (grey, n=5), 5-8min of 50μM ZD7288 (teal, n=5), 20 min Control solution (black, n =5), or 20 min 50μM ZD7288 (blue, n=5). Input resistance significantly increased after 20 min 50μM ZD7288 incubation compared to control. Analysis performed with Mann Whitney U test (see methods). * p < 0.05. n.s., not significant.

**Figure 4 – figure supplement 1.**
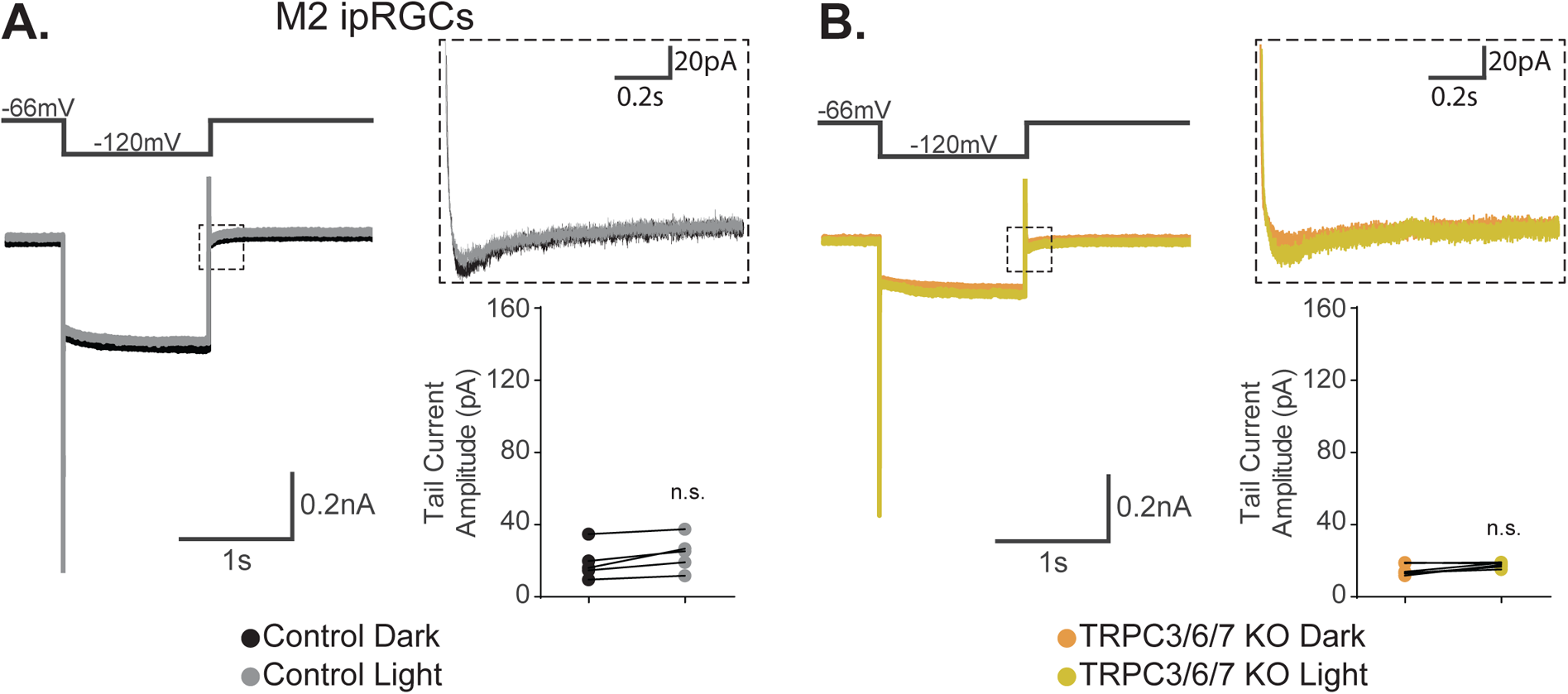
M2 HCN tail current is not modulated by light. (A) WT M2 ipRGC in dark (black) hyperpolarized from −66mV to −120mV and stepped back to the original holding potential. Cell was then placed in 90s of 480 nm light (6.08 × 10^15^ photons · cm^−2^ · s^−1^) (light grey) and subjected to the same voltage clamp protocol (gray). Tail currents are boxed and expanded in inset. Absolute value of the tail current amplitude is plotted for M2 cells in dark (black, n=5) and in light (gray, n=5). There is no significant change in the tail current amplitude in light. Recordings made in presence of synaptic blockers. (B) TRPC3/6/7 KO M2 ipRGC in dark (orange) hyperpolarized from −66mV to −120mV and stepped back to the original holding potential. Cell was then placed in 90s of 480 nm light (6.08 × 10^15^ photons · cm^−2^ · s^−1^) (yellow) and subjected to the same voltage clamp protocol. Tail currents are boxed and expanded in inset. Absolute value of the tail current amplitude is plotted for TRPC3/6/7 KO M2 cells in dark (orange, n=6) and in light (yellow, n=6). There is no significant change in the tail current amplitude in light. Recordings made in presence of synaptic blockers. Performed statistical analysis with Wilcoxon signed-rank test (see methods). n.s., not significant.

**Figure 4 – figure supplement 2.**
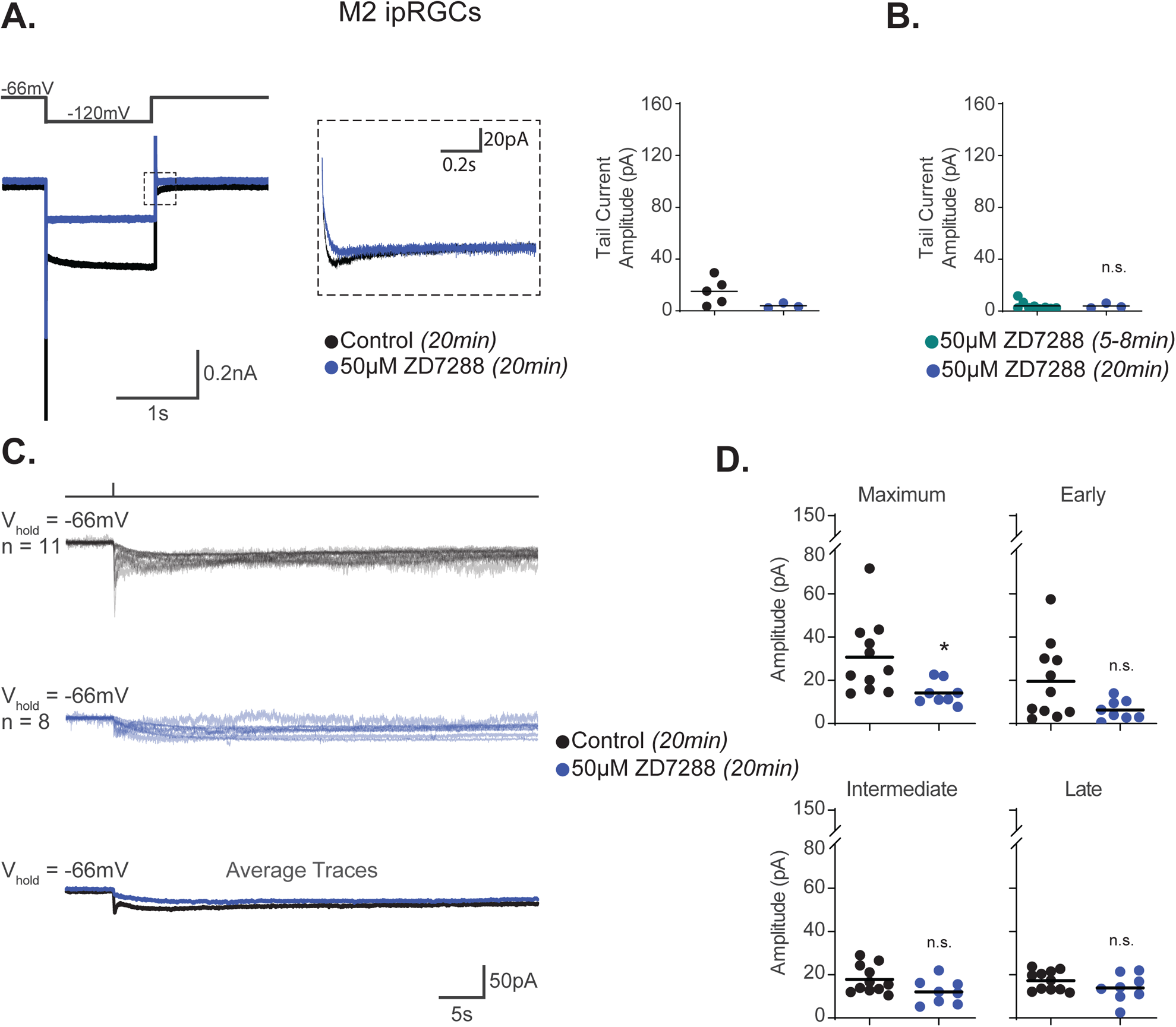
Prolonged application of ZD7288 reduces the M2 photocurrent via off-target effects. (A) WT M2 ipRGC is hyperpolarized from −66mV to −120mV and stepped back to the original holding potential after 20 min in control solution (black, n=5) or after 20 min incubation 50μM ZD7288 (blue, n=3). Tail currents are boxed and expanded in inset. Absolute value of the tail current amplitude is plotted for each cell. (B) Absolute value of HCN tail current amplitudes are similar after 5-8 (teal, n=9) vs. 20 min incubation with 50μM ZD7288 (blue, n=3), indicating no further blockade of HCN channels with longer incubation. (C) M2 photocurrent recorded after 20 min in control solution (black, n=11) or after 20 min incubation with 50μM ZD7288 (blue, n=8). Bottom row, overlaid average light response trace for each group. (D) Maximum, Early, Intermediate, and Late photocurrent amplitudes of M2 cells in 20min Control solution 20min (black, n=11) of 50μM ZD7288 (blue, n=8). The absolute value of the Maximum (p=0.0409) amplitude of M2 cells incubated in 50μM ZD7288 for 20min is significantly reduced. The Early, Intermediate, and Late photocurrent amplitudes were reduced but are not statistically significant. The results show a partial reduction of the M2 photocurrent as reported by Jiang et al., 2018. Analysis performed with Mann Whitney U test (see methods). * p < 0.05. n.s., not significant.* p < 0.05. n.s., not significant.

**Figure 5 – figure supplement 1.**
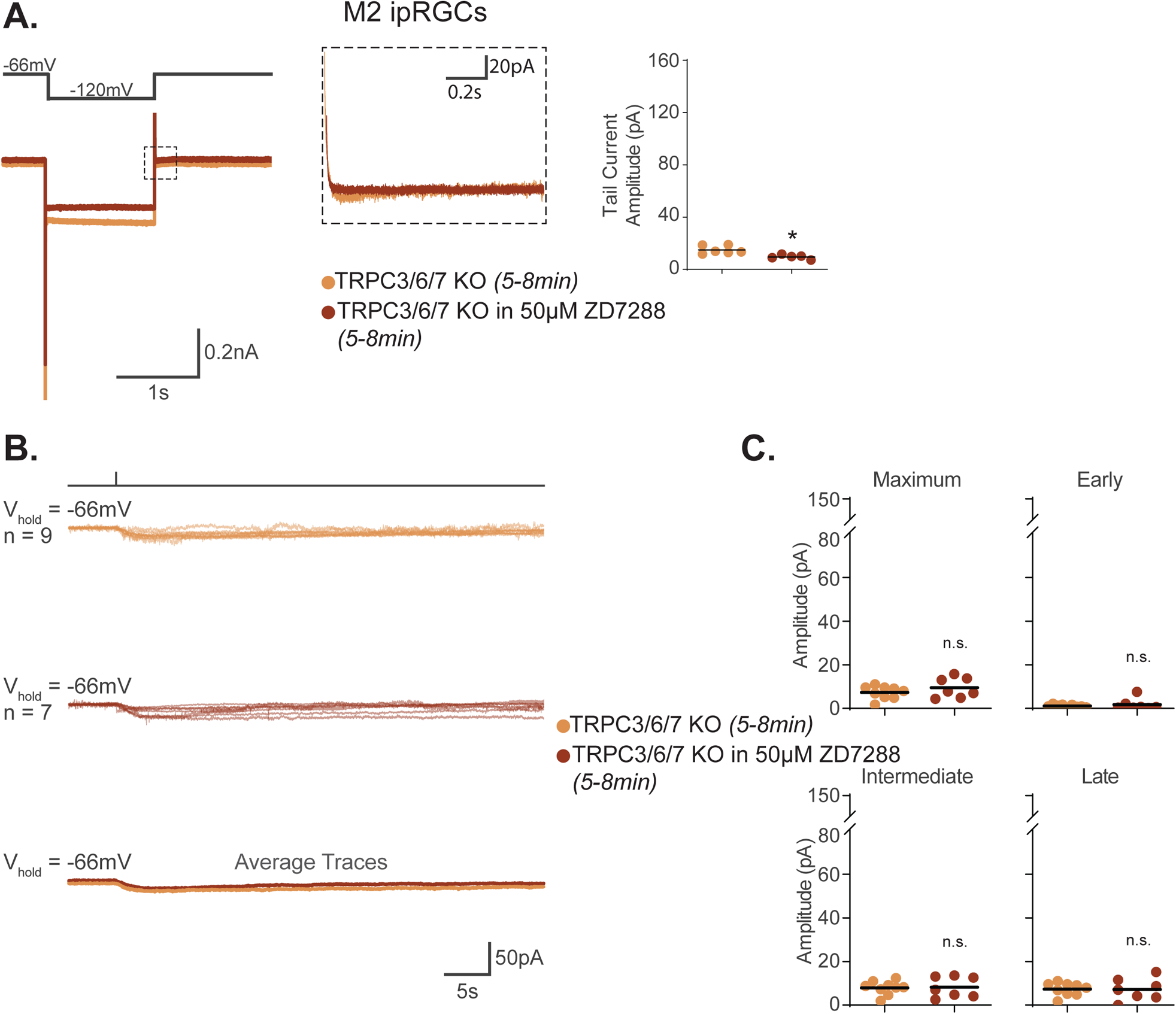
Blockade of HCN channels does not reduce the TRPC3/6/7 KO M2 photocurrent. (A) TRPC3/6/7 KO M2 ipRGC (orange), hyperpolarized from −66mV to −120mV and stepped back to the original holding potential. TRPC3/6/7 KO M2 cell is incubated in 50μM ZD7288 for 5-8min (maroon) and subjected to the same voltage protocol. Tail currents are boxed and magnified in inset. Right, top row, magnified boxed tail currents. Absolute value of the HCN tail current amplitude for TRPC3/6/7 KO M2 cells (orange, n=6) and cells incubated in 50μM ZD7288 for 5-8min (maroon, n=5). 50μM ZD7288 for 5-8min fully blocked HCN mediated tail currents in M2 ipRGCs lacking TRPC3/6/7 channels (p=0.0043). (B) Individual light responses of TRPC3/6/7 KO M2 cells (orange, n=9) or cells in 50μM ZD7288 for 5-8min (maroon, n=7). Bottom row shows the overlaid average light response trace for TRPC3/6/7 KO (orange) and in 50μM ZD7288 for 5-8min (maroon). (C) Absolute value of the current amplitudes for the light responses in (B) are graphed to compare the Maximum, Early, Intermediate, and Late component amplitudes of TRPC3/6/7 KO cells (orange, n=9) and cells in 50μM ZD7288 for 5-8min of (maroon, n=7). Light response of TRPC3/6/7 KO M2 cells in 50μM ZD7288 for 5-8min is unaffected despite blockade of HCN channels. Performed statistical analysis with Mann Whitney U test (see methods). * p < 0.05. n.s., not significant.

**Figure 6 – figure supplement 1.**
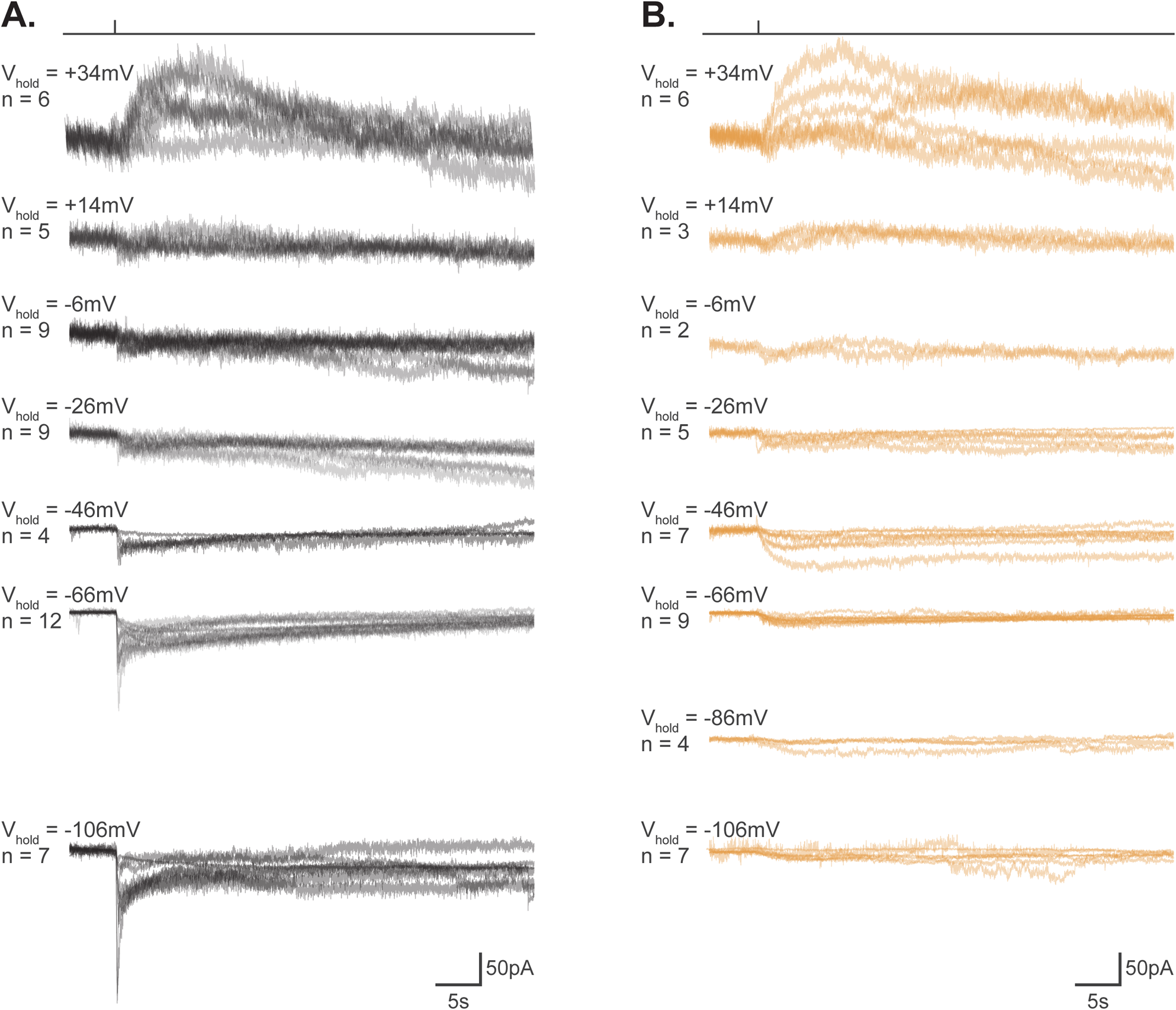
Individual M2 photocurrents used to construct the I-V relationship. (A-B) M2 photocurrent for individual Control (black) or TRPC3/6/7 KO M2 cells at various holding potentials (−106 mV to +34 mV). For each holding potential, 4-12 Control cells were recorded for a total of n=52 cells and 2-9 TRPC3/6/7 KO cells were recorded for a total of n=43 cells. Cell recordings were made in Opn4-GFP retinas in the synaptic blockers and presented with a flash of blue (480nm) light (6.08 × 10^15^ photons · cm^−2^ · s^−1^).

**Figure 9 – figure supplement 1.**
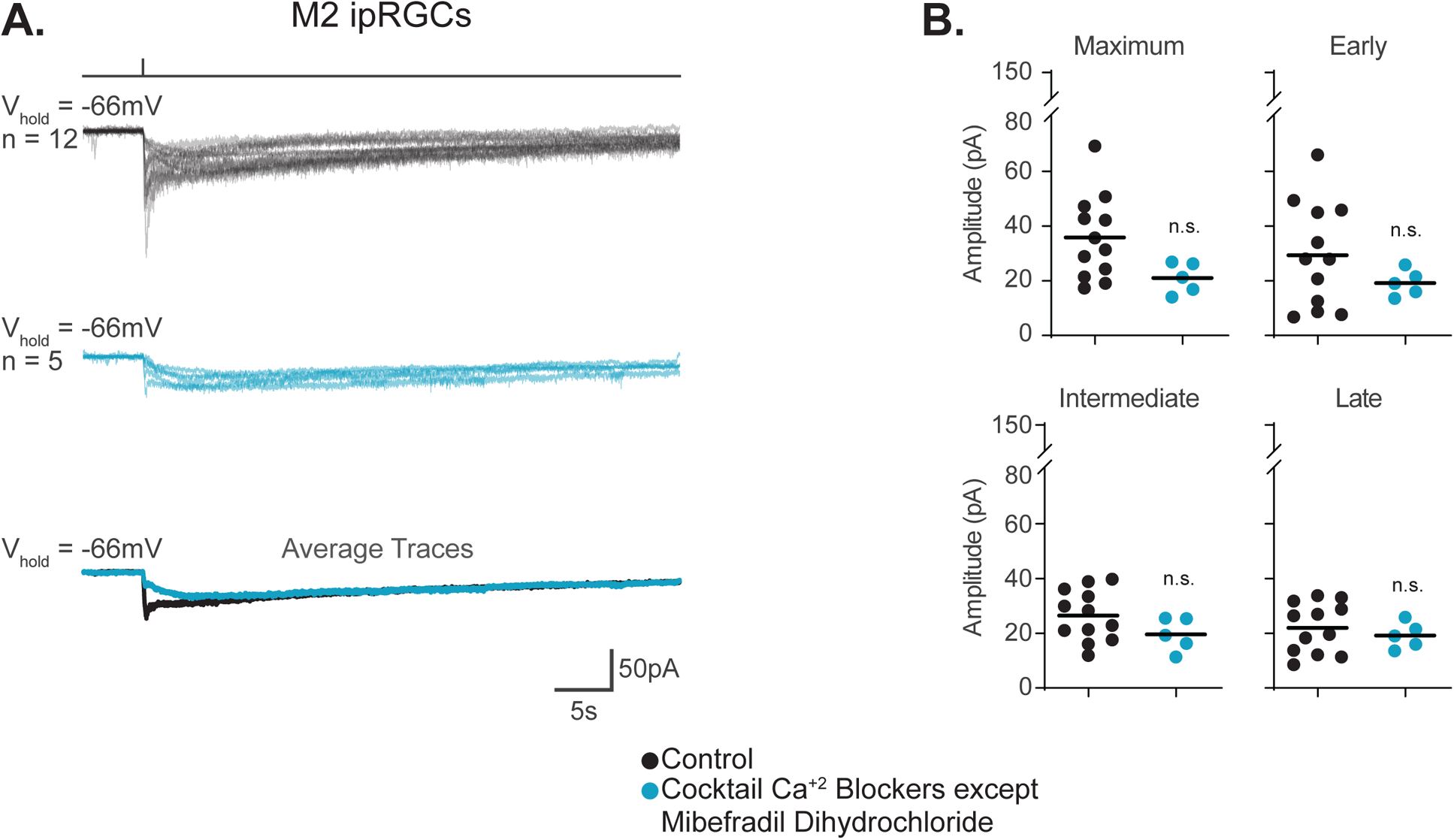
Blockade of non-T-type VGCCs does not reduce the M2 photocurrent. (A) Photocurrent of M2 ipRGCs in Control solution containing synaptic blockers alone (black, n=12) or in a Cocktail of synaptic blockers and VGCC blockers except for 10μM Mibefradil Dihydrochloride (sky blue, n=5). Blockers used: 10μM nifedipine, 5μM nimodipine, 400nM ω-agatoxin IVA, 3μM ω-conotoxin GVIA, and 3nM SNX-482. Bottom row are the overlaid average light response traces. (B) Absolute value of photocurrent amplitudes for cells in (A). Analysis performed with Mann Whitney U test (see methods). n.s., not significant.

